# Behavioral performance requirements for division of labor influence adaptive brain mosaicism in a socially complex ant

**DOI:** 10.1101/2021.07.03.450997

**Authors:** I.B. Muratore, E.M. Fandozzi, J.F.A. Traniello

**Author notes:** Corresponding author., 1-516-477-7714, 5 Cummington Mall, Boston 02215, MA, USA.

## Abstract

Brain evolution is hypothesized to be driven by neuroarchitectural requirements for behavioral performance. Assessments of such needs should be informed by the nature of sensory and motor processes underpinning behavior. We developed a novel metric to estimate the relative neuroanatomical investments required to perform tasks varying in sensorimotor and processing demands across polymorphic and polyethic workers of the leafcutter ant *Atta cephalotes* and quantified brain size and structure to examine their correspondence with our computational approximations. Investment in multi-sensory integration and motor requirements for task performance was estimated to be greatest for media workers whose leaf-harvesting repertoire involves the most diverse and demanding sensory and motor processes, including plant discrimination, leaf cutting, and fragment transportation. Volumetric analysis of confocal brain images revealed that absolute brain size increased with worker size and compartmental scaling allometries among functionally specialized brain compartments differed among polymorphic workers. The mushroom bodies, centers of sensory integration and learning, and the antennal lobes, which process olfactory inputs, were significantly larger in medias than in minim workers (fungal gardeners) and major workers (“soldiers”), which had lower estimated task-related neural demands. Minims had a proportionally larger central complex, perhaps to control navigation in subterranean fungal garden chambers. These results indicate that variation in task performance requirements has selected for adaptive variation in brain size and mosaic scaling.

## Introduction

Identifying the selective forces that contribute to the evolution of brain size and patterns of investment in functionally specialized brain centers is key to understanding the organization of behavior. Social life – as an agent of selection – has been predicted to both increase (Adolphs, 2003; Dunbar, 1998, 2009) and decrease brain size (Godfrey & Gronenberg, 2019; Gronenberg & Riveros, 2009; Jaffe & Perez, 1989; O’Donnell et al., 2018; Reséndiz-Benhumea et al., 2021; Riveros et al., 2012; Sulger et al., 2014), and/or alter compartmental scaling relationships (DeCasien & Higham, 2019; Muscedere & Traniello 2012; O’Donnell et al., 2015; O’Donnell et al., 2018; Smaers & Soligo, 2013). Factors favoring enlargement of the brain are constantly debated (Aiello & Wheeler, 1995; DeCasien et al., 2017; Lihoreau et al., 2012; Navarrete et al., 2016; Wartel et al., 2019). Assessments of sensory, motor, and processing demands for behavioral performance are required to make predictions about brain size and adaptive scaling given the cost of neural tissue, but the nature and extent of such demands are rarely estimated in analyzes that link behavior and neuroanatomy.

Insects have been proposed as models to examine behavioral and/or cognitive evolution (Lihoreau et al., 2019; Simons & Tibbetts, 2019), and some clades may facilitate the quantification of behavioral demands required to understand the evolution of neural systems (Boogert et al., 2018; Muratore & Traniello, 2020). Ants, as exemplars of insect sociality, form complex societies that may be characterized by striking patterns of task specialization associated with ergonomic division of labor (Beshers & Fewell, 2001; Hölldobler & Wilson, 1990) and worker size-related variation in innate task routines and/or cognitive capabilities. The brains of workers are composed of functionally specialized compartments involved in visual and olfactory processing (optic lobes [OL] and antennal lobes [AL], respectively), higher-order processing, learning, and memory (mushroom bodies [MB]), navigation, orientation, and movement (central complex [CX], also associated with the MBs) (Currier et al., 2020; Green et al., 2019; Kamhi et al., 2020; Le Moël et al., 2019; Pisokas et al., 2020; Sun et al., 2020), and mandibular control and gustation (the suboesophageal zone [SEZ]). The remainder of the central brain (ROCB) is composed of several protocerebral compartments thought to integrate sensory information with the CX (Currier et al., 2020; Green et al., 2019; Strausfeld, 2012). Although brain size may be limited as a predictor of computational power and cognitive capacity (Chittka & Niven, 2009; Muscedere et al., 2014), the volumes and structural elaboration of visual and olfactory neuropils and the MBs in ants are considered to correlate with processing capability and have ethological significance (Farris, 2011; Gronenberg, 2001). Brain size and compartmental allometries (Gordon et al., 2017; Muscedere & Traniello, 2012) and covariance among brain compartments (Ilieş et al., 2015; Kamhi et al., 2019) correlate with variability in worker size and task performance. Task experience affects MB size (Durst et al., 1994; Fahrbach, 2006; Gronenberg et al., 1996), and large MBs may be associated with increased behavioral flexibility (Gronenberg & Riveros, 2009; O’Donnell et al., 2015; Riveros et al., 2012). Complexity in colony organization may therefore involve selection for either smaller, neurally differentiated worker brains (Feinerman & Traniello, 2015; Gronenberg & Riveros, 2009; Lihoreau et al., 2012; O’Donnell et al., 2018; O’Donnell et al., 2015; Riveros et al., 2012; Sulger et al., 2014) or larger brains (Wehner et al., 2007) potentially able to metabolically offset increased production and operation costs (Kamhi et al., 2016).

Social organization in fungus-growing ants is characterized by morphologically differentiated workers in derived species (Hölldobler & Wilson, 2010; Muratore & Traniello, 2020). Colonies of leafcutter ants, *Atta,* have strongly polymorphic workers categorized into size-based groups (subcastes) differing in labor roles and efficiencies of task performance (Figure 1) (Wilson, 1980a, 1980b). Worker size-related specializations include fungal care, nursing immatures, leaf selection, cutting, and transport, waste management and hygienic behavior, and colony defense. These tasks differ in requirements for multimodal sensory perception, integrative processing, and motor functions associated with monitoring the growth of fungus and brood development, decoding plant chemistry, leaf harvesting, navigation, recognizing infectious agents, detecting environmental hazards, and combating predators and parasites (Arenas & Roces, 2016a, 2016b, 2017; Arganda et al., 2020; Goes et al., 2020; Green & Kooij, 2018; Groh et al., 2014; Howard et al., 1996; Saverschek & Roces, 2011). This pattern of division of labor characterizes *Atta cephalotes*, whose workers range in size from 0.5 to more than 4.5mm in head width and are divisible into groups according to task repertoires (Table 1). Brain size and structural variation could reflect differences in behavioral diversity: workers displaying the broadest task repertoire and corresponding behavioral and/or cognitive demands could require higher investment in neural tissue. To examine factors contributing to brain evolution in *A. cephalotes*, we evaluated the distribution of functional needs for sensory perception, integration, and sensorimotor functions associated with task performance across polymorphic workers. We determined patterns of variation in behavioral demands associated with task performance and work environments to provide a semiquantitative estimate of brain tissue requirements to efficaciously perform tasks.

**Figure 1.**
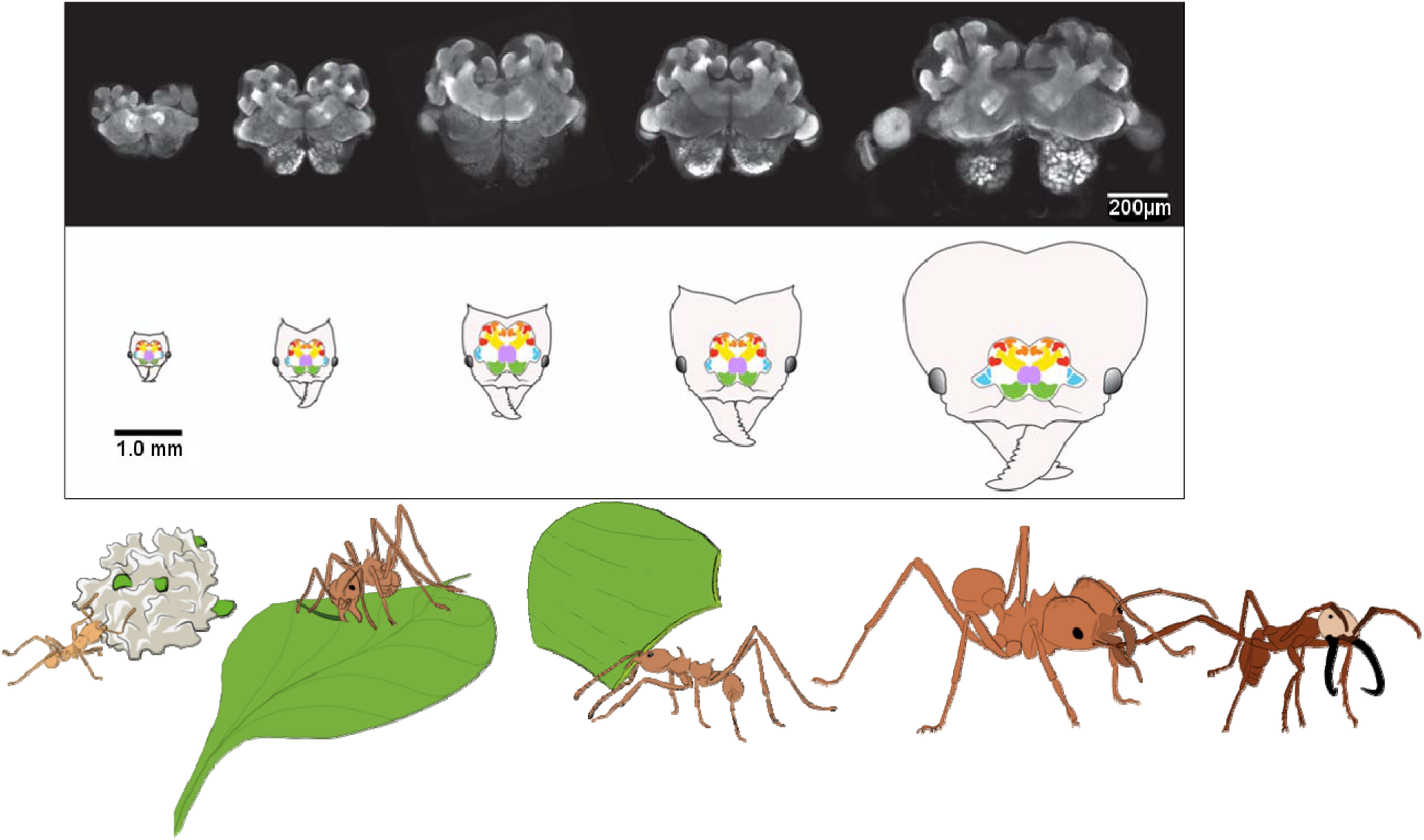
Confocal images of *A. cephalotes* polymorphic worker brains (top row). Structural diagrams of brains illustrating color-coded neuropils whose volumes were measured (middle row; compartments not drawn to scale). Blue = OL, green = AL, orange = MB-MC, red = MB-LC, yellow = MB-P, purple = SEZ, pink = CX. Diagram depicting task performance in relation to worker size (bottom row; minim, two images of medias, major attacking antagonistic species).

**Table 1.**
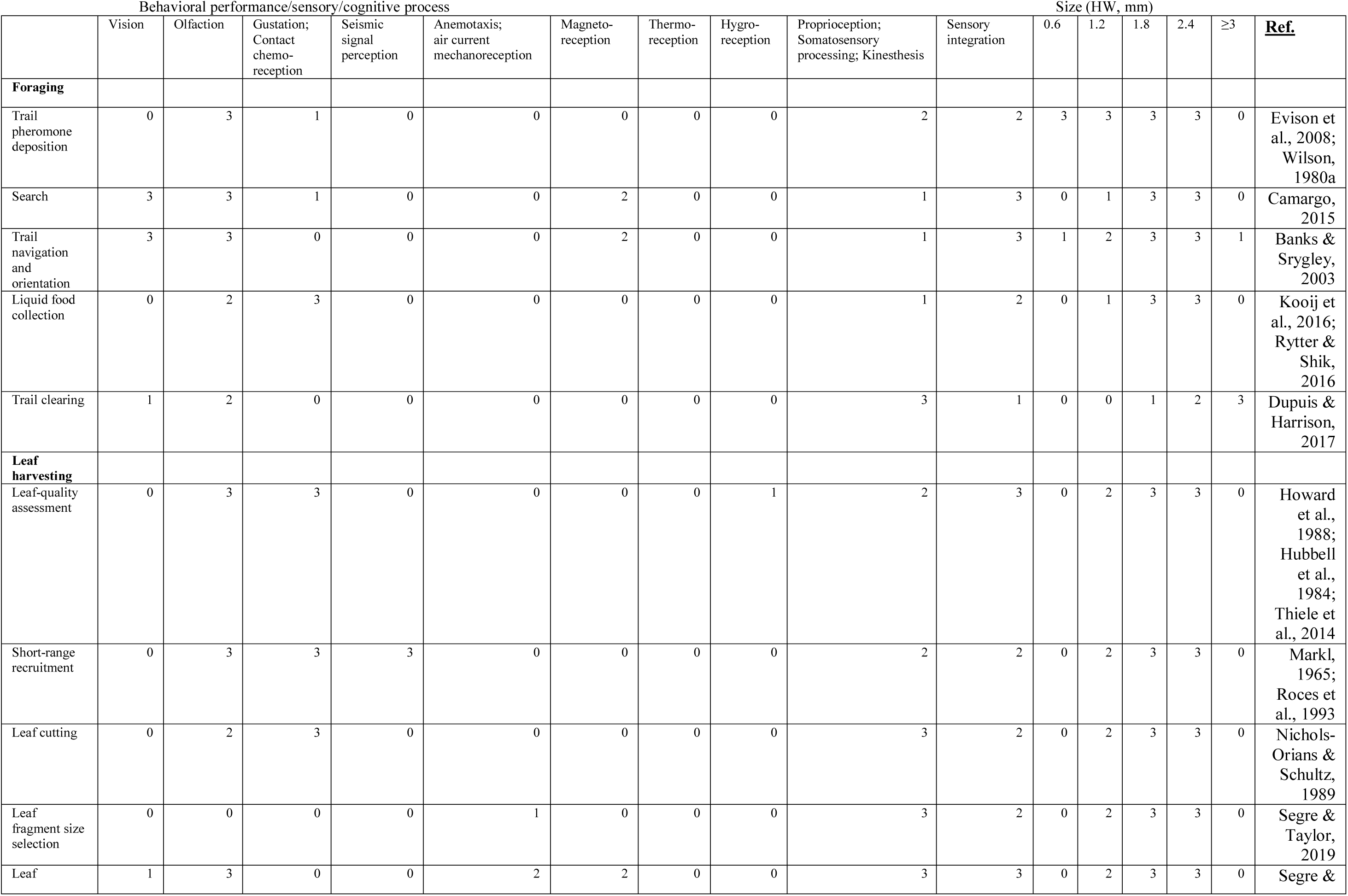
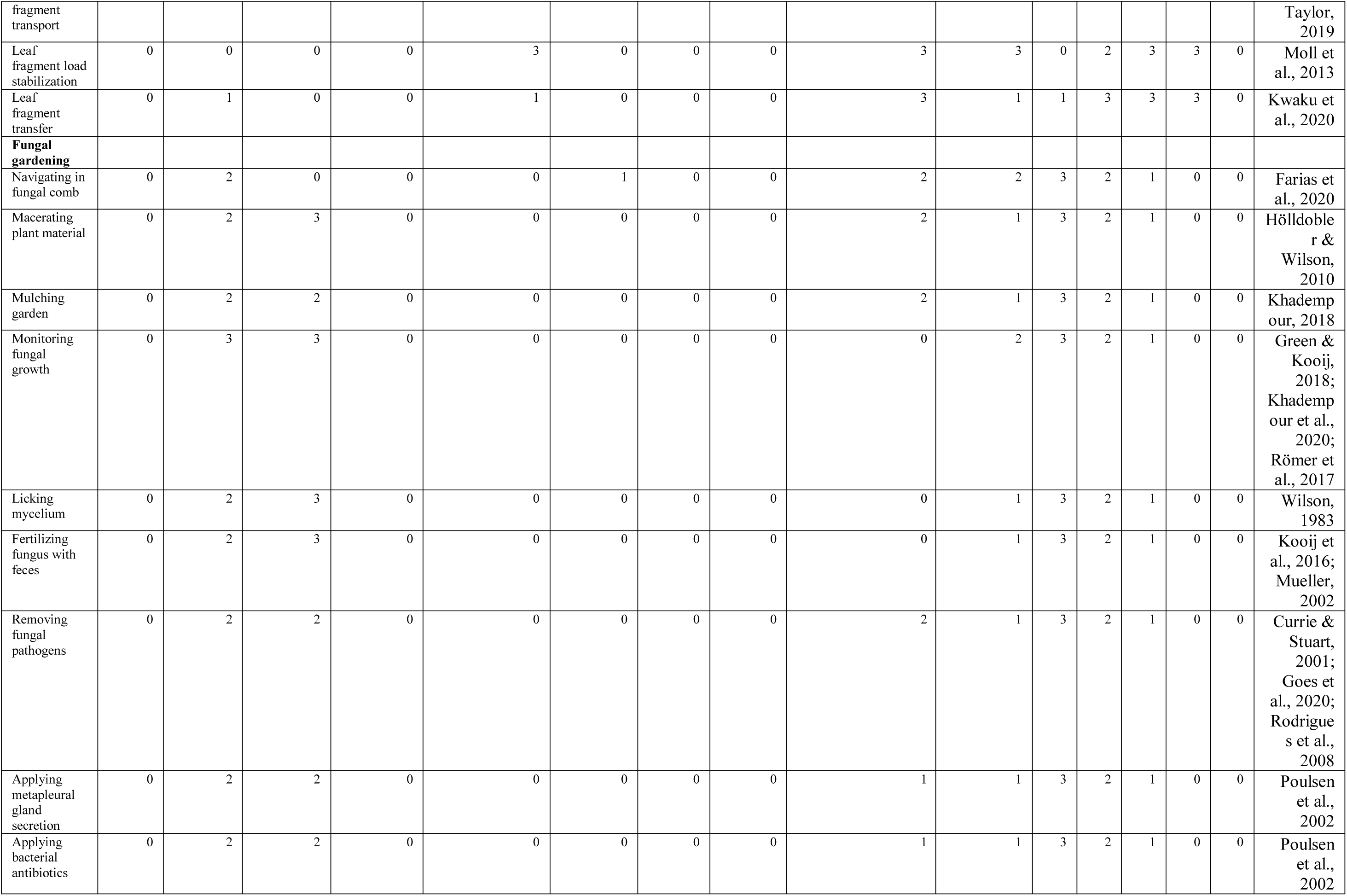
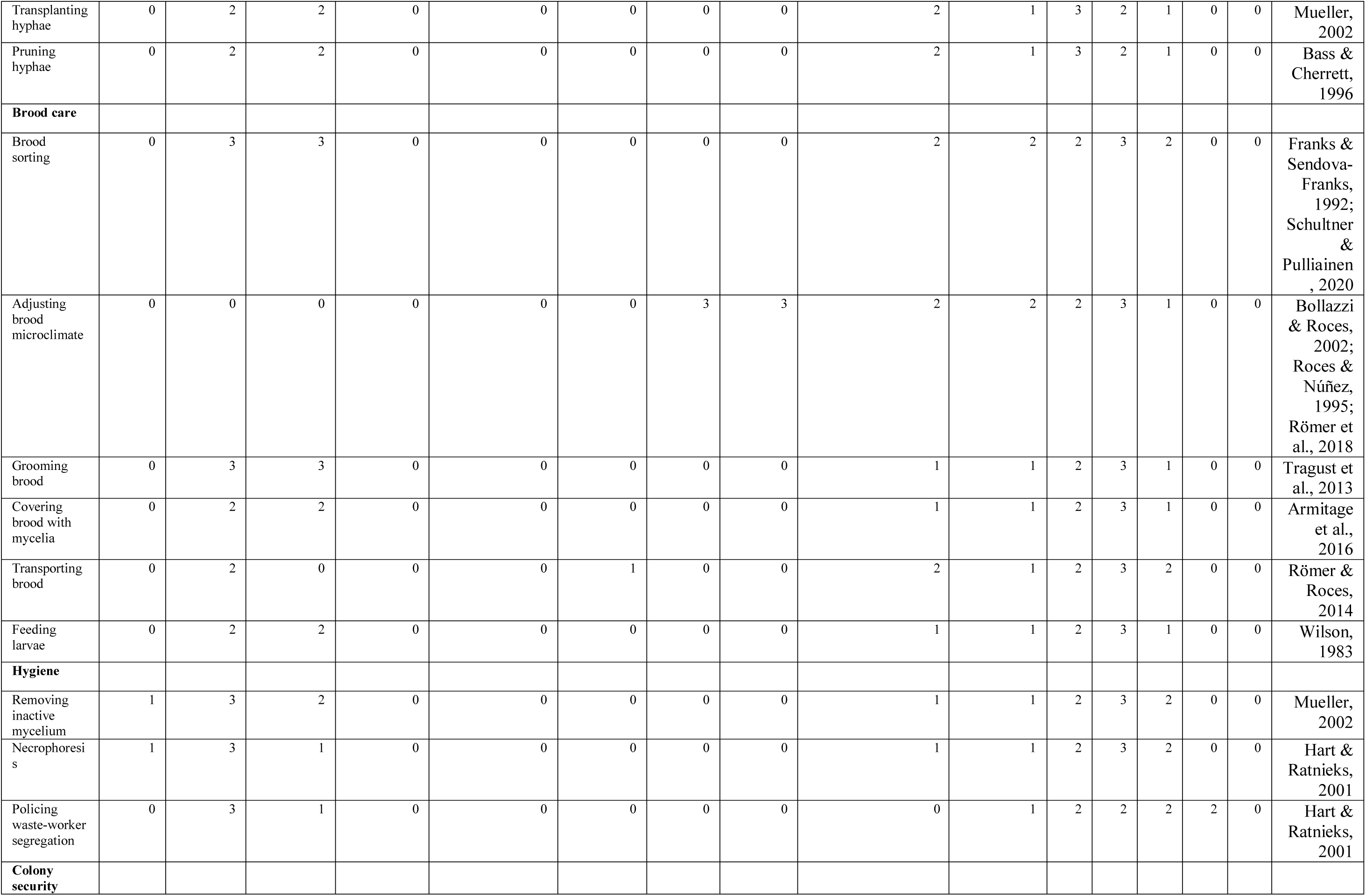
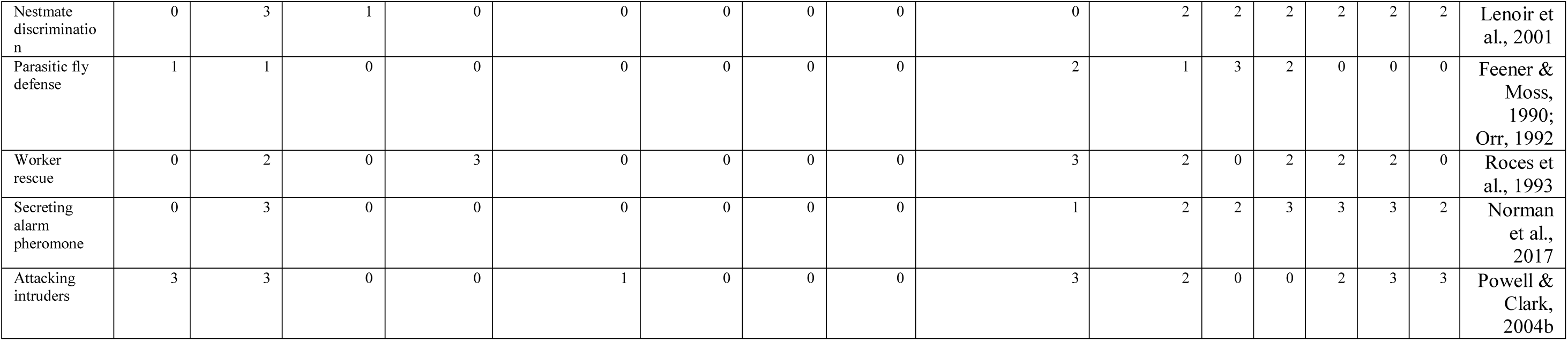
Tasks performed by *A. cephalotes* workers, their relative reliance on sensory modalities, sensory input processing, high-order integration, and motor coordination in relation to division of labor. Sensory and sensorimotor inputs of tasks and processing are rated on a 0-3 scale depending on the role an input or process is considered to serve in a given task. 0 = not involved, 1 = possible role, 2 = likely role, and 3 = significant role. The same scale was used to rate the performance frequency of a given task in the repertoire of different-size workers. The “proprioception, somatosensory processing and kinesthesis” category refers to requirements above common body and appendage motor control systems across polymorphic workers that reflect greater kinesthetic coordination than standing, walking without load carriage, feeding, mouthpart and appendage movement, self-grooming, and allogrooming. Value assignments are based on published results and personal observations.

| Behavioral performance/sensory/cognitive process |  |  |  |  |  |  |  |  |  |  | Size (HW, mm) |  |  |  |  |  |
| --- | --- | --- | --- | --- | --- | --- | --- | --- | --- | --- | --- | --- | --- | --- | --- | --- |
|  | Vision | Olfaction | Gustation;<br>Contact<br>chemo-<br>reception | Seismic<br>signal<br>perception | Anemotaxis;<br>air current<br>mechanoreception | Magneto-<br>reception | Thermo-<br>reception | Hygro-<br>reception | Proprioception;<br>Somatosensory<br>processing; Kinesthesia | Sensory<br>integration | 0.6 | 1.2 | 1.8 | 2.4 | ≥3 | <u>Ref.</u> |
| Foraging |  |  |  |  |  |  |  |  |  |  |  |  |  |  |  |  |
| Trail<br>pheromone<br>deposition | 0 | 3 | 1 | 0 | 0 | 0 | 0 | 0 | 2 | 2 | 3 | 3 | 3 | 3 | 0 | Evison et<br>al., 2008;<br>Wilson,<br>1980a |
| Search | 3 | 3 | 1 | 0 | 0 | 2 | 0 | 0 | 1 | 3 | 0 | 1 | 3 | 3 | 0 | Camargo,<br>2015 |
| Trail<br>navigation<br>and<br>orientation | 3 | 3 | 0 | 0 | 0 | 2 | 0 | 0 | 1 | 3 | 1 | 2 | 3 | 3 | 1 | Banks &<br>Srygley,<br>2003 |
| Liquid food<br>collection | 0 | 2 | 3 | 0 | 0 | 0 | 0 | 0 | 1 | 2 | 0 | 1 | 3 | 3 | 0 | Kooij et<br>al., 2016;<br>Rytter &<br>Shik,<br>2016 |
| Trail clearing | 1 | 2 | 0 | 0 | 0 | 0 | 0 | 0 | 3 | 1 | 0 | 0 | 1 | 2 | 3 | Dupuis &<br>Harrison,<br>2017 |
| Leaf<br>harvesting |  |  |  |  |  |  |  |  |  |  |  |  |  |  |  |  |
| Leaf-quality<br>assessment | 0 | 3 | 3 | 0 | 0 | 0 | 0 | 1 | 2 | 3 | 0 | 2 | 3 | 3 | 0 | Howard<br>et al.,<br>1988;<br>Hubbell<br>et al.,<br>1984;<br>Thiele et<br>al., 2014 |
| Short-range<br>recruitment | 0 | 3 | 3 | 3 | 0 | 0 | 0 | 0 | 2 | 2 | 0 | 2 | 3 | 3 | 0 | Markl,<br>1965;<br>Roces et<br>al., 1993 |
| Leaf cutting | 0 | 2 | 3 | 0 | 0 | 0 | 0 | 0 | 3 | 2 | 0 | 2 | 3 | 3 | 0 | Nichols-Orians &<br>Schultz,<br>1989 |
| Leaf<br>fragment size<br>selection | 0 | 0 | 0 | 0 | 1 | 0 | 0 | 0 | 3 | 2 | 0 | 2 | 3 | 3 | 0 | Segre &<br>Taylor,<br>2019 |
| Leaf | 1 | 3 | 0 | 0 | 2 | 2 | 0 | 0 | 3 | 3 | 0 | 2 | 3 | 3 | 0 | Segre & |
| fragment transport |  |  |  |  |  |  |  |  |  |  |  |  |  |  |  | Taylor, 2019 |
| Leaf fragment load stabilization | 0 | 0 | 0 | 0 | 3 | 0 | 0 | 0 | 3 | 3 | 0 | 2 | 3 | 3 | 0 | Moll et al., 2013 |
| Leaf fragment transfer | 0 | 1 | 0 | 0 | 1 | 0 | 0 | 0 | 3 | 1 | 1 | 3 | 3 | 3 | 0 | Kwaku et al., 2020 |
| **Fungal gardening** |  |  |  |  |  |  |  |  |  |  |  |  |  |  |  |  |
| Navigating in fungal comb | 0 | 2 | 0 | 0 | 0 | 1 | 0 | 0 | 2 | 2 | 3 | 2 | 1 | 0 | 0 | Farias et al., 2020 |
| Macerating plant material | 0 | 2 | 3 | 0 | 0 | 0 | 0 | 0 | 2 | 1 | 3 | 2 | 1 | 0 | 0 | Hölldobler & Wilson, 2010 |
| Mulching garden | 0 | 2 | 2 | 0 | 0 | 0 | 0 | 0 | 2 | 1 | 3 | 2 | 1 | 0 | 0 | Khadempour, 2018 |
| Monitoring fungal growth | 0 | 3 | 3 | 0 | 0 | 0 | 0 | 0 | 0 | 2 | 3 | 2 | 1 | 0 | 0 | Green & Kooij, 2018; Khadempour et al., 2020; Römer et al., 2017 |
| Licking mycelium | 0 | 2 | 3 | 0 | 0 | 0 | 0 | 0 | 0 | 1 | 3 | 2 | 1 | 0 | 0 | Wilson, 1983 |
| Fertilizing fungus with feces | 0 | 2 | 3 | 0 | 0 | 0 | 0 | 0 | 0 | 1 | 3 | 2 | 1 | 0 | 0 | Kooij et al., 2016; Mueller, 2002 |
| Removing fungal pathogens | 0 | 2 | 2 | 0 | 0 | 0 | 0 | 0 | 2 | 1 | 3 | 2 | 1 | 0 | 0 | Currie & Stuart, 2001; Goes et al., 2020; Rodrigues et al., 2008 |
| Applying metapleural gland secretion | 0 | 2 | 2 | 0 | 0 | 0 | 0 | 0 | 1 | 1 | 3 | 2 | 1 | 0 | 0 | Poulsen et al., 2002 |
| Applying bacterial antibiotics | 0 | 2 | 2 | 0 | 0 | 0 | 0 | 0 | 1 | 1 | 3 | 2 | 1 | 0 | 0 | Poulsen et al., 2002 |
| Transplanting hyphae | 0 | 2 | 2 | 0 | 0 | 0 | 0 | 0 | 2 | 1 | 3 | 2 | 1 | 0 | 0 | Mueller, 2002 |
| Pruning hyphae | 0 | 2 | 2 | 0 | 0 | 0 | 0 | 0 | 2 | 1 | 3 | 2 | 1 | 0 | 0 | Bass & Cherrett, 1996 |
| **Brood care** |  |  |  |  |  |  |  |  |  |  |  |  |  |  |  |  |
| Brood sorting | 0 | 3 | 3 | 0 | 0 | 0 | 0 | 0 | 2 | 2 | 2 | 3 | 2 | 0 | 0 | Franks & Sendova-Franks, 1992; Schultner & Pulliainen, 2020 |
| Adjusting brood microclimate | 0 | 0 | 0 | 0 | 0 | 0 | 3 | 3 | 2 | 2 | 2 | 3 | 1 | 0 | 0 | Bollazzi & Roces, 2002; Roces & Núñez, 1995; Römer et al., 2018 |
| Grooming brood | 0 | 3 | 3 | 0 | 0 | 0 | 0 | 0 | 1 | 1 | 2 | 3 | 1 | 0 | 0 | Tragust et al., 2013 |
| Covering brood with mycelia | 0 | 2 | 2 | 0 | 0 | 0 | 0 | 0 | 1 | 1 | 2 | 3 | 1 | 0 | 0 | Armitage et al., 2016 |
| Transporting brood | 0 | 2 | 0 | 0 | 0 | 1 | 0 | 0 | 2 | 1 | 2 | 3 | 2 | 0 | 0 | Römer & Roces, 2014 |
| Feeding larvae | 0 | 2 | 2 | 0 | 0 | 0 | 0 | 0 | 1 | 1 | 2 | 3 | 1 | 0 | 0 | Wilson, 1983 |
| **Hygiene** |  |  |  |  |  |  |  |  |  |  |  |  |  |  |  |  |
| Removing inactive mycelium | 1 | 3 | 2 | 0 | 0 | 0 | 0 | 0 | 1 | 1 | 2 | 3 | 2 | 0 | 0 | Mueller, 2002 |
| Necrophoresis | 1 | 3 | 1 | 0 | 0 | 0 | 0 | 0 | 1 | 1 | 2 | 3 | 2 | 0 | 0 | Hart & Ratnieks, 2001 |
| Policing waste-worker segregation | 0 | 3 | 1 | 0 | 0 | 0 | 0 | 0 | 0 | 1 | 2 | 2 | 2 | 2 | 0 | Hart & Ratnieks, 2001 |
| **Colony security** |  |  |  |  |  |  |  |  |  |  |  |  |  |  |  |  |
| Nestmate discrimination | 0 | 3 | 1 | 0 | 0 | 0 | 0 | 0 | 0 | 2 | 2 | 2 | 2 | 2 | 2 | Lenoir et al., 2001 |
| Parasitic fly defense | 1 | 1 | 0 | 0 | 0 | 0 | 0 | 0 | 2 | 1 | 3 | 2 | 0 | 0 | 0 | Feener & Moss, 1990; Orr, 1992 |
| Worker rescue | 0 | 2 | 0 | 3 | 0 | 0 | 0 | 0 | 3 | 2 | 0 | 2 | 2 | 2 | 0 | Roces et al., 1993 |
| Secreting alarm pheromone | 0 | 3 | 0 | 0 | 0 | 0 | 0 | 0 | 1 | 2 | 2 | 3 | 3 | 3 | 2 | Norman et al., 2017 |
| Attacking intruders | 3 | 3 | 0 | 0 | 1 | 0 | 0 | 0 | 3 | 2 | 0 | 0 | 2 | 3 | 3 | Powell & Clark, 2004b |

To identify how brain size, compartmental scaling, and differential task performance demands of division of labor are associated and thus determine correspondence with our estimate, we quantified patterns of sensory, higher-order processing, and motor neuropil investment. This allowed us to link variation in task repertoires and their sensory challenges among workers to size scaling among functionally differentiated brain compartments. Based on an assessment of worker size-related sensory and motor functions, olfaction needs, and higher-order processing, we hypothesized that intermediate-size (media, leaf-harvesting) workers would have higher MB investment than small (minims, fungus-gardening) or large (majors, defensive) workers. We additionally hypothesized that our estimate would accurately describe the pattern of investment in the ALs, with this neuropil proportionally largest in medias due to the need for processing diverse olfactory cues. We also determined the fit of our estimates to the proportional volumes of the CX and OLs.

## Methods

### Colony collection and culturing

Incipient *A. cephalotes* colonies were collected in Trinidad during 2016. Colonies (Ac09, Ac16, Ac20, Ac21) were cultured in a Harris environmental chamber under a 12-hour light: 12-hour dark cycle at 55% humidity and 25°C at Boston University, and Costa Rican colonies (M1 and M2) were cultured in an environmental chamber under a 12-hour light: 12-hour dark cycle at 20°C at the Boston Museum of Science. All colonies were housed in large plastic bins (30cm x 46cm x 28cm), whose floors served as a foraging arena and for waste disposal. Smaller plastic boxes (11cm x 18cm x 13cm) interconnected by plastic tubes (1cm diameter) served as chambers for the fungus. Colonies were provisioned with washed and pesticide-free leaves from rhododendron, rose, lilac, andromeda, bramble, oak, sugar maple, willow, and beech trees (as available), and organic baby spinach, romaine, arugula, frisée, and oat flakes.

### Behavioral performance demands and estimates of required neuroanatomical support

We integrated fungus-growing ant adaptive morphology and behavior (Hölldobler & Wilson, 2010; Wilson, 1980a, 1980b) with personal observations and data from the literature (references in Table 1) to inform our estimate of needs for sensory integration motor control in *A. cephalotes* worker behavior. Based on our results of research on visual system evolution in *A. cephalotes* (Arganda et al., 2020), we assumed greater investment in neuropils such as the MBs and ALs would be required to process more diverse stimulus arrays and coordinate sensorimotor processes. For example, tasks such as leaf selection, cutting, and transport involve olfactory discrimination, proprioception and mechanosensory and muscular systems to control the mandibles, appendages, head position, and direction of movement (Currier et al., 2020; Green et al., 2019; Khalife et al., 2018), whereas other tasks differ substantially in these needs. Estimates for task performance frequency were based on published results of studies of worker size-related behavior (references listed in Table 1) validated by unpublished data (Muratore et al., in prep.). Sensory integration and other cognitive process requirements were based on overlap in the known sensory capacities of ants and documented instances of behaviors being disrupted through manipulations of the brain or sensory pathways (see references in Table 1).

Approximations of neuropil requirements for sensory integration were calculated as a sum of each worker group-task performance process combination from estimates in Table 1 according to the following equation:

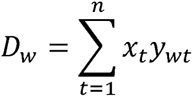

*D_W_* = Estimated requirement for neuronal substrate to process and integrate sensory and/or sensorimotor inputs for a given worker size group

*x_t_ =* Sensory integration-task demand score, the estimated degree of sensory integration required to perform a given task *t*

*y_wt_ =* Worker size group task performance score, estimating the tendency of a given worker group w to perform a given task *t*

*n* = total number of tasks

This equation integrates contributions from the type of brain investments likely to be necessary for the performance of individual tasks with the frequency with which different types of workers are likely to perform them in order to generate a hypothesis about the likelihood of selection acting to prioritize or deprioritize sensory integration in the brains of different workers. *x_t_* was rated on a 0-3 scale according to the role an input or process is thought to serve in a specific task. 0 = not involved, 1 = possible role, 2 = likely role, and 3 = significant role. *y_wt_* was rated on the same scale according to the likelihood of a given worker size group to perform a specific task.

### Immunohistochemistry and confocal microscopy

Mature fully sclerotized workers collected from colonies Ac09, Ac16, Ac20, and Ac21 were decapitated immediately prior to brain dissection and fixation. Workers were sampled from five worker size groups identified by head width (HW): minims (0.6mm±0.1mm), medias (1.2mm±0.1mm, 1.8±0.1mm, or 2.4mm±0.1mm), and majors (3.0mm or larger). Brains (n=30) from workers sampled from Ac09, Ac20, and Ac21 were dissected in ice-cold HEPES Buffered Saline (HBS), placed in 16% zinc-formaldehyde (Ott, 2008) and fixed overnight at room temperature (RT) on a shaker. Whole brains were processed to visualize the presynaptic protein synapsin. Fixed brains were washed in HBS six times, 10 minutes per wash, and fixed in Dent’s Fixative (80% MeOH, 20% DMSO) for minimally 1 hour. Brains were then washed in 100% methanol and either stored at −17°C or immediately processed. Brains were washed in 0.1M Tris buffer (pH=7.4) and blocked in PBSTN (5% neutral goat serum, 0.005% sodium azide in 0.2% PBST) at RT for 1 hour before incubation for 3 days at RT in primary antibody (1:30 SYNORF 1 in PBSTN; monoclonal antibody anti-synorf 3C11 obtained from DSHB, University of Iowa, IA, USA; 62). They were washed 6×10 minutes in 0.2% PBST and incubated in the secondary antibody (1:100 AlexaFluor 488 goat anti-mouse in PBSTN) for 4 days at RT. Brains were then washed a final time (6×10 minutes in 0.2% PBST) and dehydrated in an ethanol and PBS series (10 minutes per concentration, 30/50/70/95/100/100% ethanol in 1x PBS), then cleared with and immersed in methyl salicylate, and mounted on stainless steel glass windowed slides for imaging.

Brains were imaged with a Nikon C2 confocal microscope and images were manually annotated using Amira 6.0 software to quantify neuropil volumes (not including cell bodies). We recorded the volumes of OL, AL, MB, CX, SEZ, and ROCB. We also measured substructures of the MB: the medial calyxes (MB-MC), lateral calyxes (MB-LC), and peduncle and lobes (MB-P). These were combined to quantify total MB size (MB-S) across worker size groups. For bilateral structures, one hemisphere was measured, and for compartments located along the brain midline (SEZ and CX), the whole structure was measured (Supplementary Table 1; Supplementary Table 2).

### Volumetric analysis

Statistical *post hoc* comparisons among worker size groups and brain compartment metrics were performed using R (version 3.6.2). Absolute volumes of total measured brain volume, total brain volume scaled to head width, all individual brain compartments, and two normalized brain compartments were not normally distributed (Supplementary Table 3), and the significance of differences among groups was therefore determined using Kruskal-Wallis tests followed by Wilcoxon rank sum *post hoc* pairwise comparison with Bonferroni correction. Relative volumes were calculated by dividing the volume of the compartment of interest by total brain volume. These data were distributed normally. If ANOVA results showed a significant effect of worker size group on the proportional size of a brain compartment, Tukey’s *post hoc* tests were performed with a Bonferroni correction for multiple comparisons to determine the significance of compartment size differences among groups.

Linear regression was used to assess correspondence between estimate values of D_w_ for each worker size group and either proportional volumes of brain compartments or total brain volume scaled to worker size (i.e., the sum of all measured neuropils/HW).

Principal component analysis (PCA) was performed on log-transformed proportional volumes using the prcomp function from the base stats package in R. Linear discriminant analysis (LDA) was performed using the lda function from the MASS package in R (Liaw & Wiener, 2001). A previously collected data set of brain volume measurements from the same species with measurements taken by a different observer was used as a training set (Supplementary Table 4).

## Results

### Division of labor and requirements for task performance

Estimated requirements for sensory integration were highest in medias (1.2mm D_w_ = 124, 1.8mm = 127, 2.4mm = 102), with the estimate for minims (0.6mm D_w_ = 82) exceeding that for majors (3mm+ D_w_ = 20) (Figure 2). A linear regression of MB-S volume against D_w_ showed a significant correspondence between our estimate and the pattern of proportional investments in the MBs (Table 2, Figure 3). Similarly, MB-P volume showed moderate but significant correspondence to D_w_ estimates. MB-MC, MB-LC, OL, AL, CB, SEZ, and ROCB volumes were also compared to D_w_ estimates, all showing very low levels of explanation of variance with mixed significance (Table 2).

**Figure 2.**
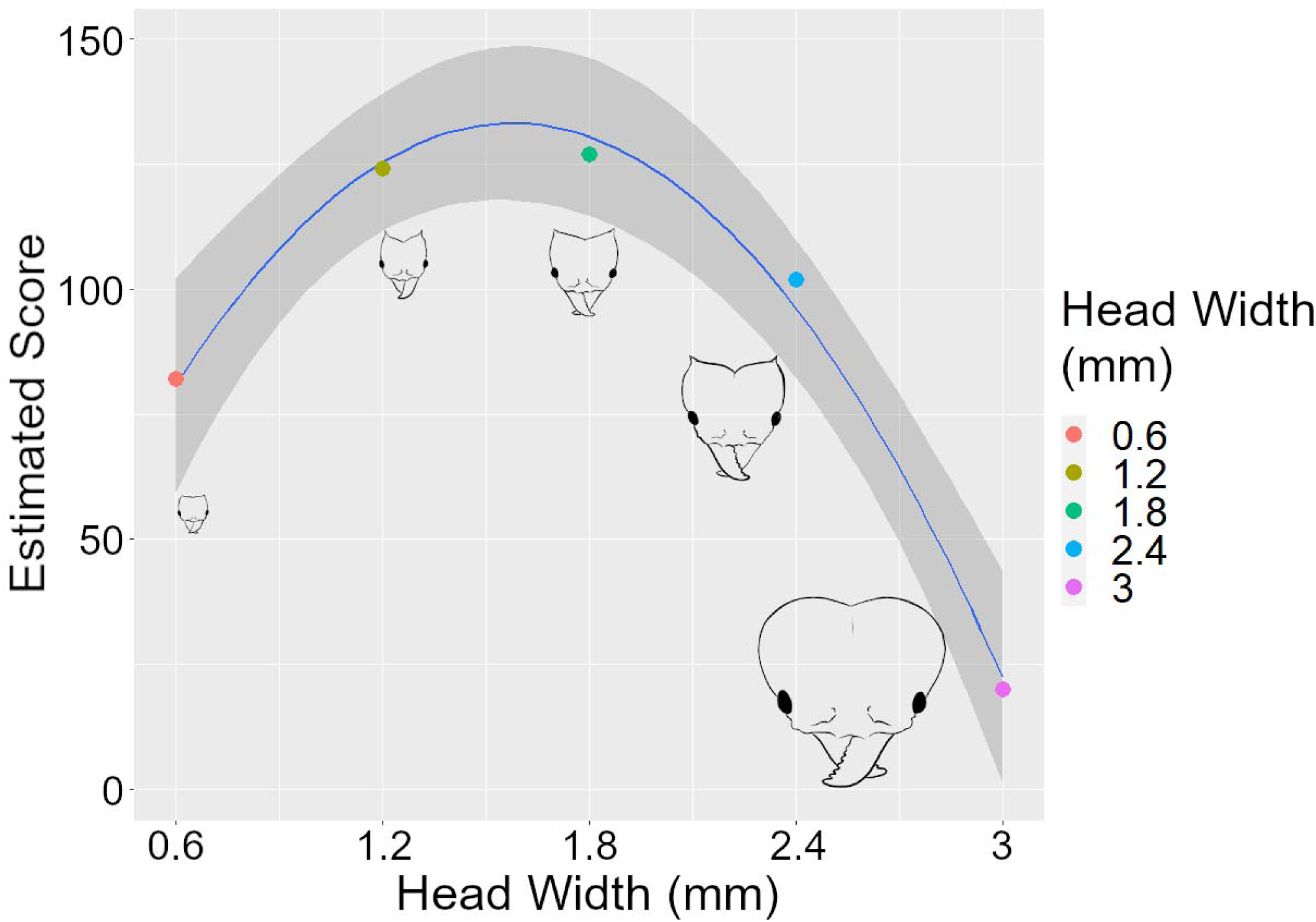
Calculated estimates (D_w_) for neuronal substrate requirements based on tasks performed by *A. cephalotes* worker size groups. Y-axis values are the sum of each sensory integration/sensorimotor function task score multiplied by the corresponding worker group-size task performance score (see Methods). Local regression curve estimates neuronal substrate requirement score as a function of HW plus HW squared (blue line) and 95% confidence interval (grey band). Heads are drawn to scale.

**Table 2.**
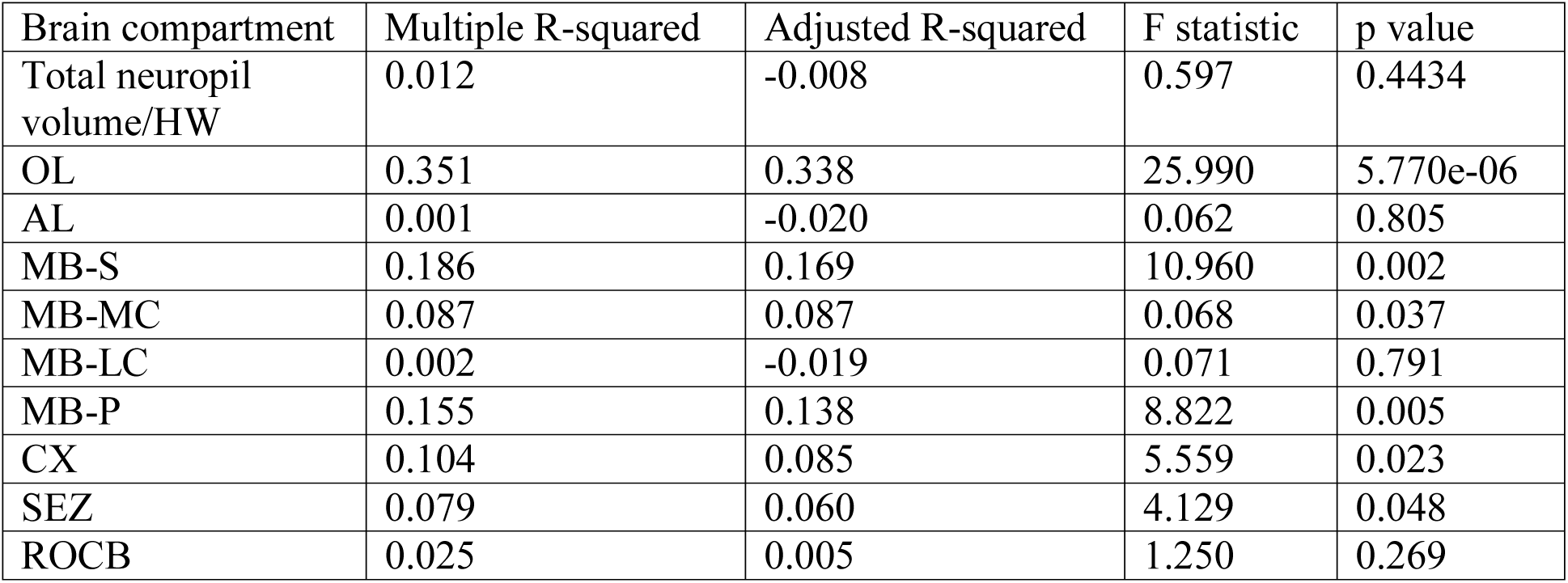
Fit of D_w_ estimates to observed patterns of proportional brain compartment investment. Linear regression statistics for the fit of a subset of categories of behavioral performance/sensory/cognitive process as predictors for the brain compartments whose function most closely corresponds to these demands. Degrees of freedom = 1, 48.

| Brain compartment | Multiple R-squared | Adjusted R-squared | F statistic | p value |
| --- | --- | --- | --- | --- |
| Total neuropil volume/HW | 0.012 | -0.008 | 0.597 | 0.4434 |
| OL | 0.351 | 0.338 | 25.990 | 5.770e-06 |
| AL | 0.001 | -0.020 | 0.062 | 0.805 |
| MB-S | 0.186 | 0.169 | 10.960 | 0.002 |
| MB-MC | 0.087 | 0.087 | 0.068 | 0.037 |
| MB-LC | 0.002 | -0.019 | 0.071 | 0.791 |
| MB-P | 0.155 | 0.138 | 8.822 | 0.005 |
| CX | 0.104 | 0.085 | 5.559 | 0.023 |
| SEZ | 0.079 | 0.060 | 4.129 | 0.048 |
| ROCB | 0.025 | 0.005 | 1.250 | 0.269 |

**Figure 3.**
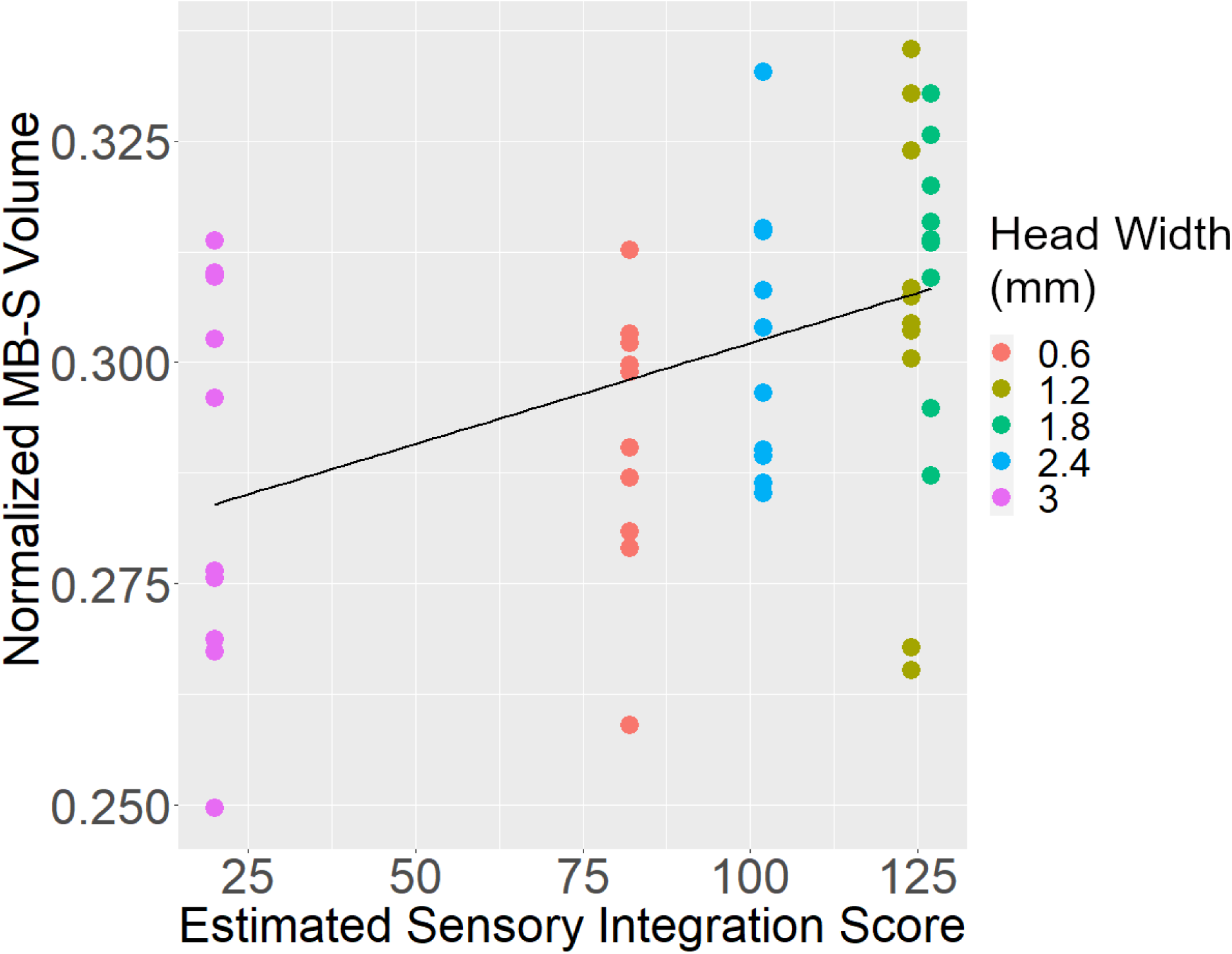
Normalized MB-S volume as a function of estimated sensory integration (Dw) with regression line. R-squared: 0.169.

### Division of labor and neural phenotypes

Absolute total brain volume of increased with worker size (Figure 4a; Table 3a; Supplementary Table 5a). Interestingly, no significant change in total brain size occurred in workers in the 0.6 to 1.8mm size range, despite a three-fold increase in body size. Only workers in the 2.4mm and 3mm+ size groups had significantly larger brains. In contrast, the smallest two size groups showed slightly but significantly larger brain volumes in relation to body size (Figure 4b; Table 3a; Supplementary Table 5b). The absolute volumes of all compartments except the CX significantly increased with worker size (Figure 5; Table 3a; Supplementary Table 6). Consistent with the pattern of total brain size, many brain compartments were significantly larger in 2.4mm or 3mm+ workers. There was a significant effect of worker size on the proportional volumes of all brain compartments except the MB-MC, MB-LC, and the SEZ (Table 3b). In contrast to the relatively uniform pattern of increase in absolute volumes, the directions of these trends differed, indicating neuroanatomical differentiation of social role phenotypes (Figure 6; Table 3b; Supplementary Table 7).

**Figure 4.**
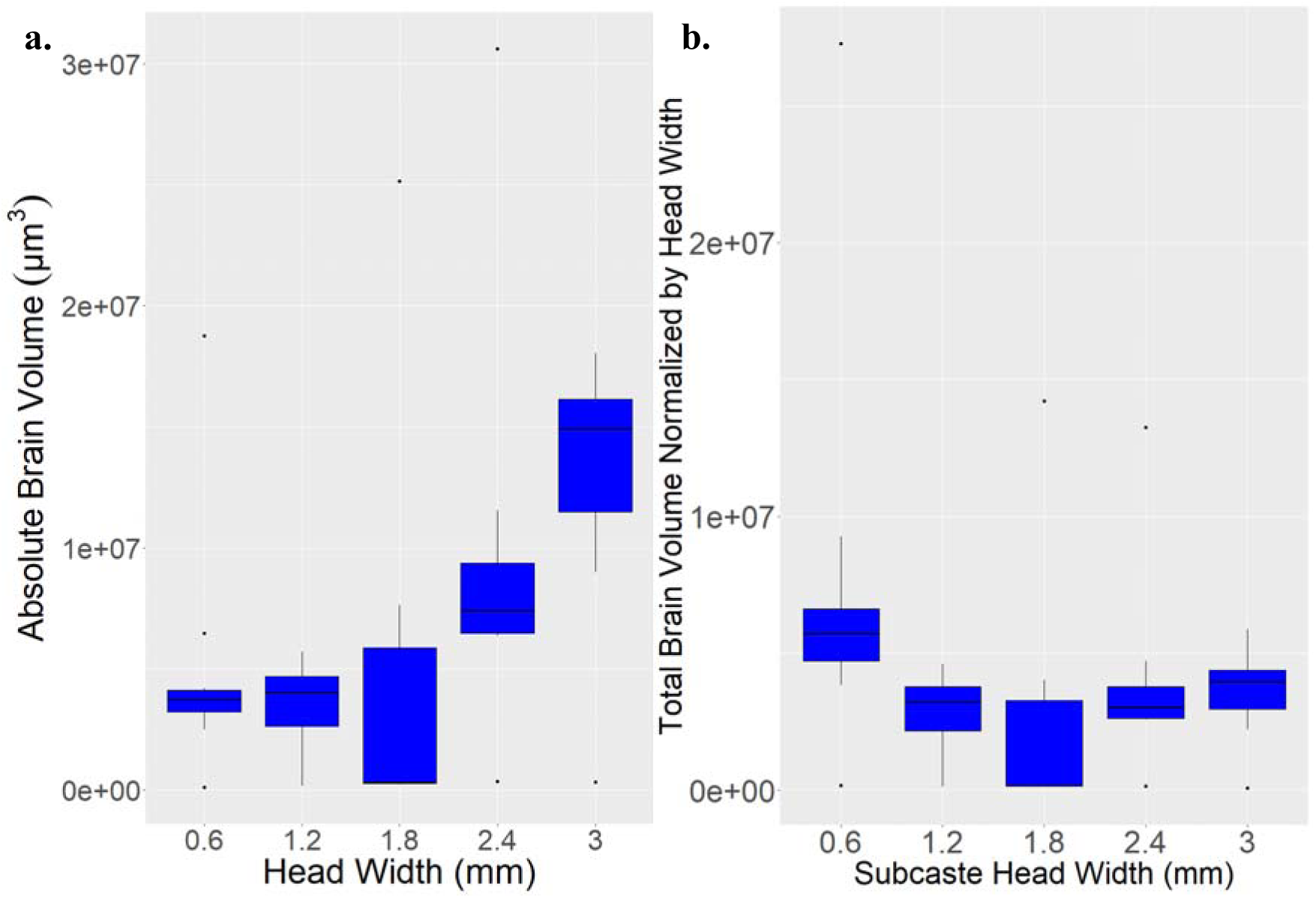
**a.** Absolute total brain volume (sum of the OLs, ALs, MB-S, CX, SEZ, and ROCB) across worker size groups (p = 0.002). **b.** Total brain volume scaled to body size (total volume/HW for each sample across worker size groups (p = 0.009). Y axis notation indicates a given number x10 raised to the power of the number indicated to the right of the e.

**Table 3.** Differences in the significance of comparisons of brain volume and brain compartment absolute **a.** and proportional volume **b.** among worker size groups. Degrees of freedom = 4.

**a.** Absolute total brain and compartmental volume
| Brain compartment | Kruskal-Wallis chi-squared | p value |
| --- | --- | --- |
| Total of neuropils | 17.291 | 0.002 |
| Total neuropil volume/HW | 13.600 | 0.009 |
| OL | 21.014 | 3.000e-04 |
| AL | 18.428 | 0.001 |
| MB-S | 17.592 | 0.002 |
| MB-MC | 17.616 | 0.002 |
| MB-LC | 17.217 | 0.002 |
| MB-P | 16.022 | 0.003 |
| CX | 5.987 | 0.200 |
| SEZ | 17.819 | 0.001 |
| ROCB | 17.048 | 0.002 |

**b.** Proportional brain compartment volumes
| Brain compartment | ANOVA F statistic | p value |
| --- | --- | --- |
| AL | 2.833 | 0.035 |
| MB-S | 66.222 | 0.013 |
| MB-MC | 1.614 | 0.187 |
| MB-LC | 1.120 | 0.359 |
| CX | 8.299 | 4.208e-05 |
| SEZ | 1.921 | 0.123 |
| ROCB | 3.233 | 0.021 |
| Brain compartment | Kruskal-Wallis chi-squared | p value |
| OL | 41.367 | 2.257e-08 |
| MB-P | 11.363 | 0.023 |

**Figure 5.**
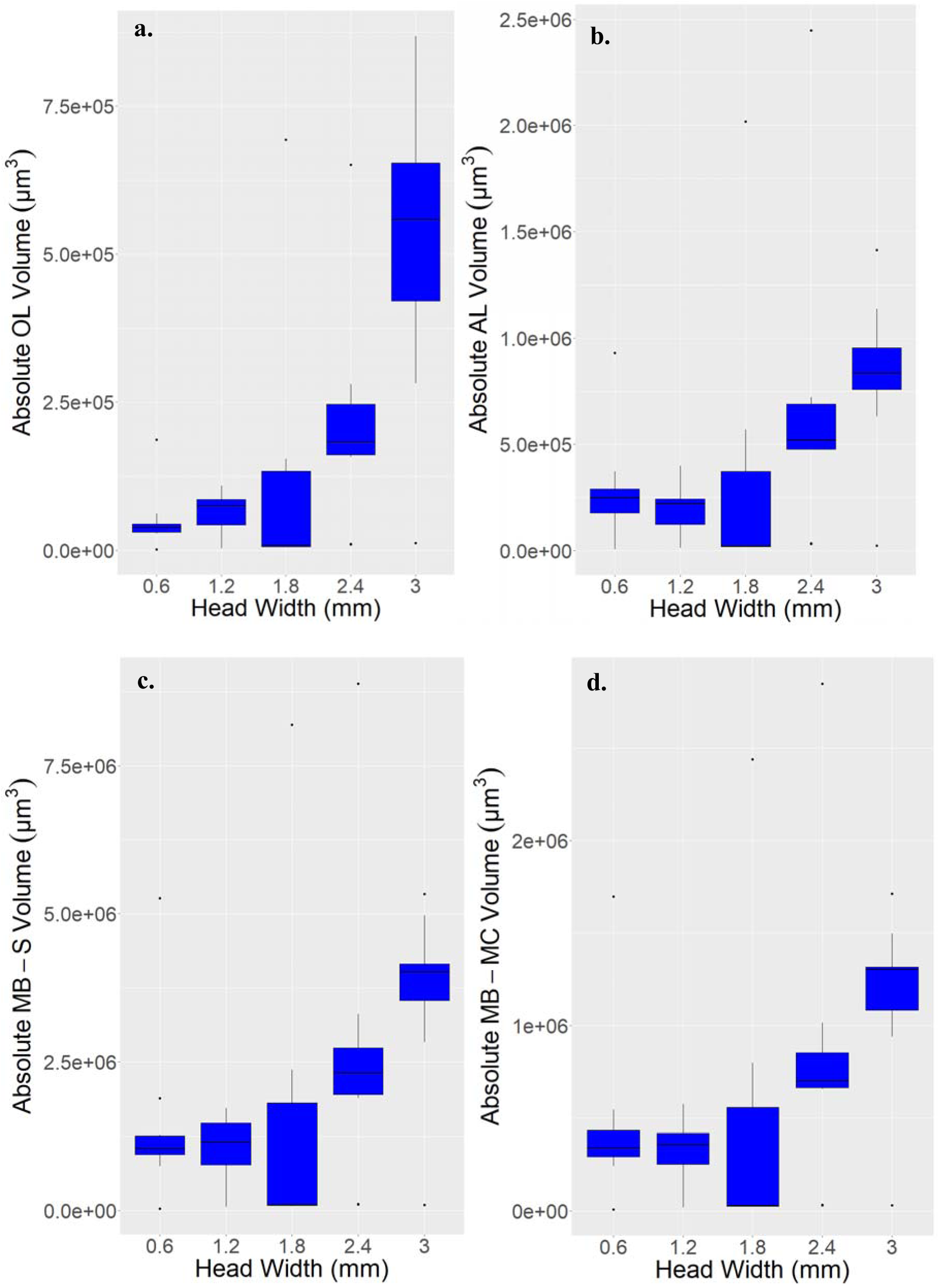
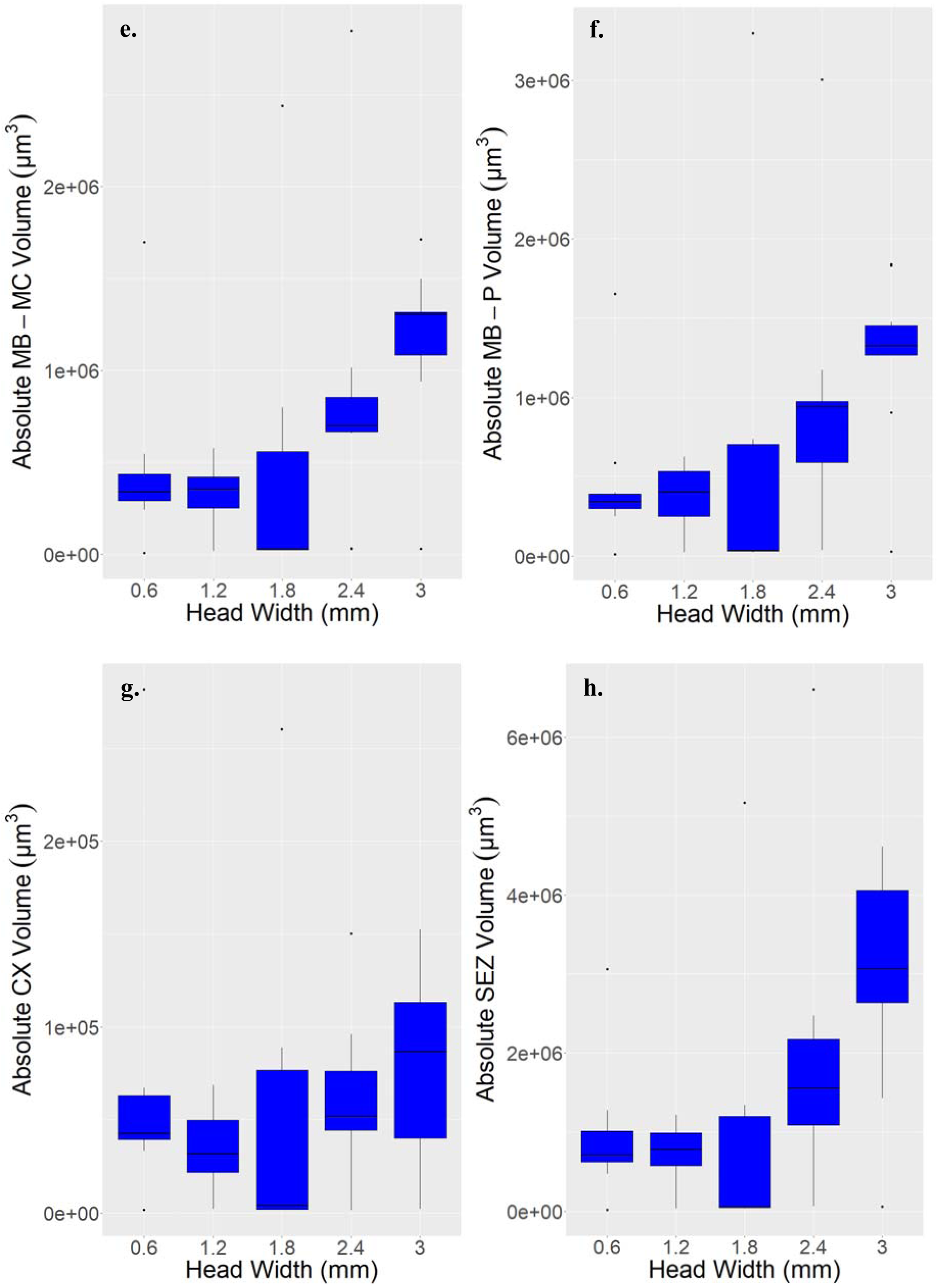
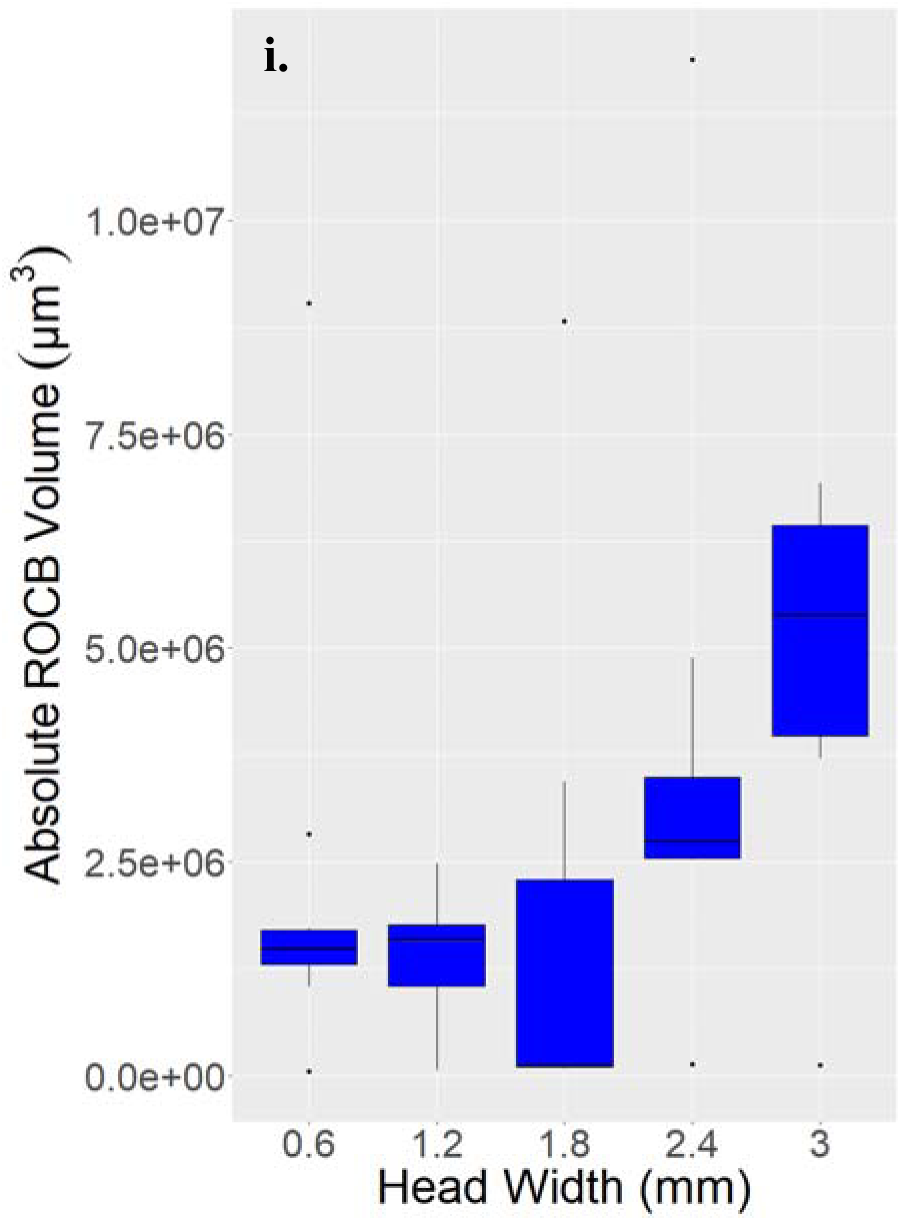
a. Absolute volume of the OL (p = 3.000e-04); b. AL (b. p = 0.001); c. MB-S (p = 0.002); d. MB-MC (p = 0.001); e. MB-LC (p = 0.002); f. MB-P (p = 0.003); g. CX (p = 0.200), h. SEZ (p = 0.001); and i. ROCB (p = 0.002) across worker size groups. Y axis notation indicates volume values x10 raised to the power of the number indicated to the right of e.

**Figure 6.**
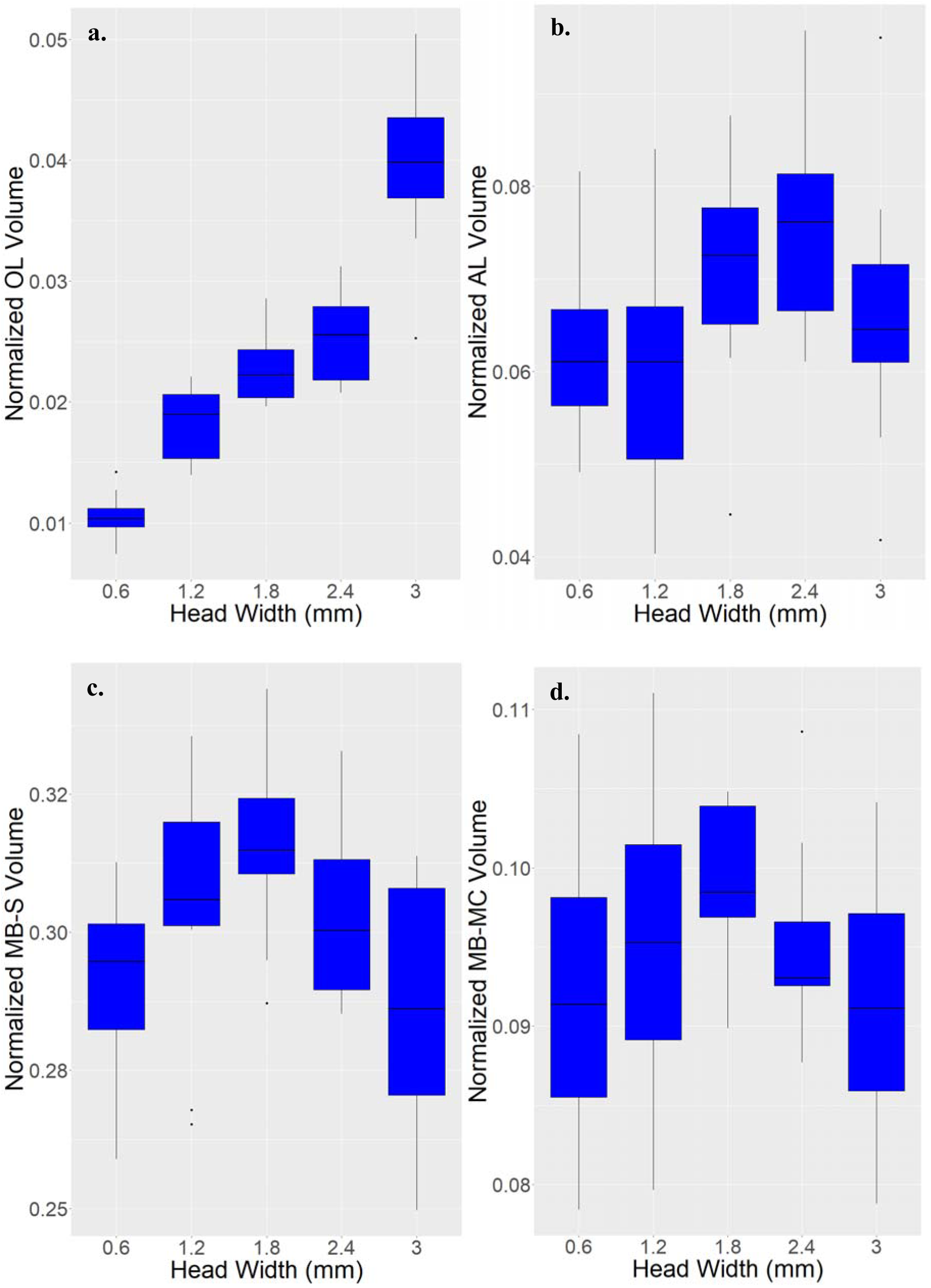
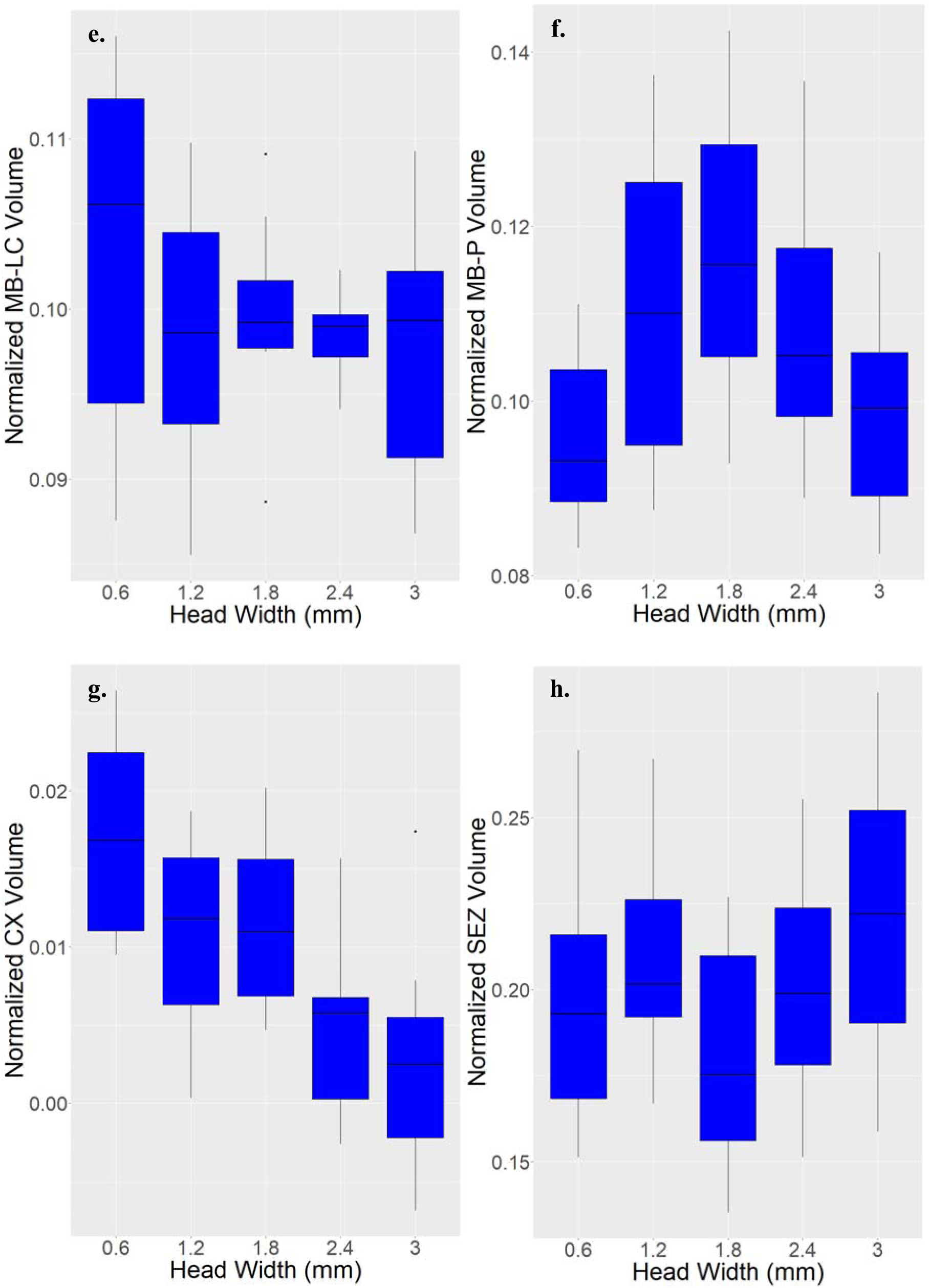
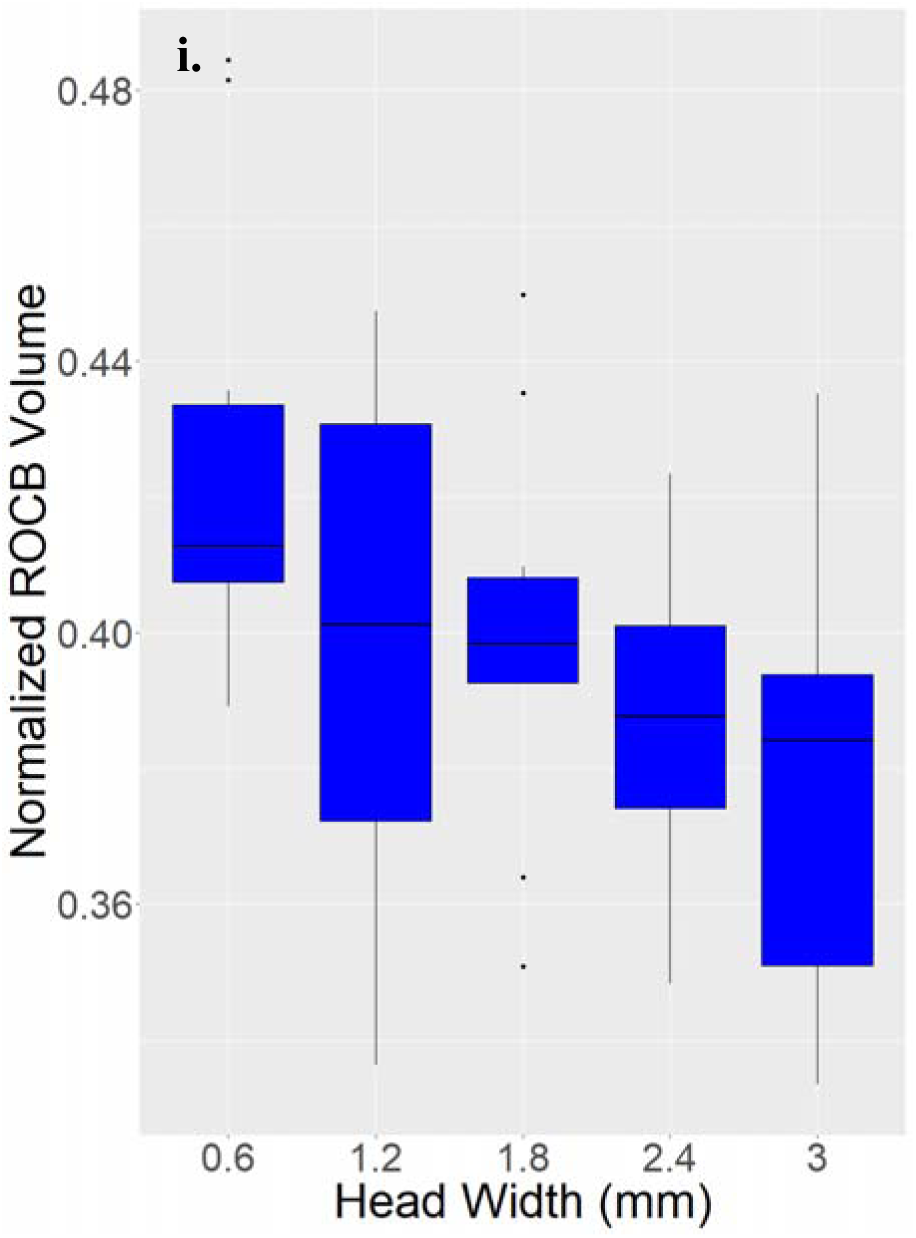
Relative investment (compartment volume as a percent of total brain volume) in the a. OL (p = 2.257e-08), AL (b., p = 0.035), MB-S (c., p = 0.013), MB-MC (d., p = 0.187), MB-LC (e., p = 0.359), MB-P (f., p = 0.023), CX (g., p = 4.208e-05), SEZ (h., p = 0.123), and ROCB (i., p = 0.021) across worker size groups. Y axis as in Figure 5.

Principal component analysis of log-transformed proportional brain volumes explained a significant portion (PC1 = 15.83%, PC = 66.04%) of the variance observed (Figure 7a). Linear discriminant analysis, using a model that included the proportional volumes of all neuropils except for the ROCB (which was colinear with other variables) and trained on a separate data set of *A. cephalotes* brain volumes, was able to classify samples in the main data set with 95.8% accuracy (Figure 7b). However, this result was found only when 1.2, 1.8, and 2.4mm worker groups were clustered as medias. LDA using five worker size groups (0.6, 1.2, 1.8, and 3mm+) and the same testing and training data sets achieved 54.2% accuracy.

**Figure 7.**
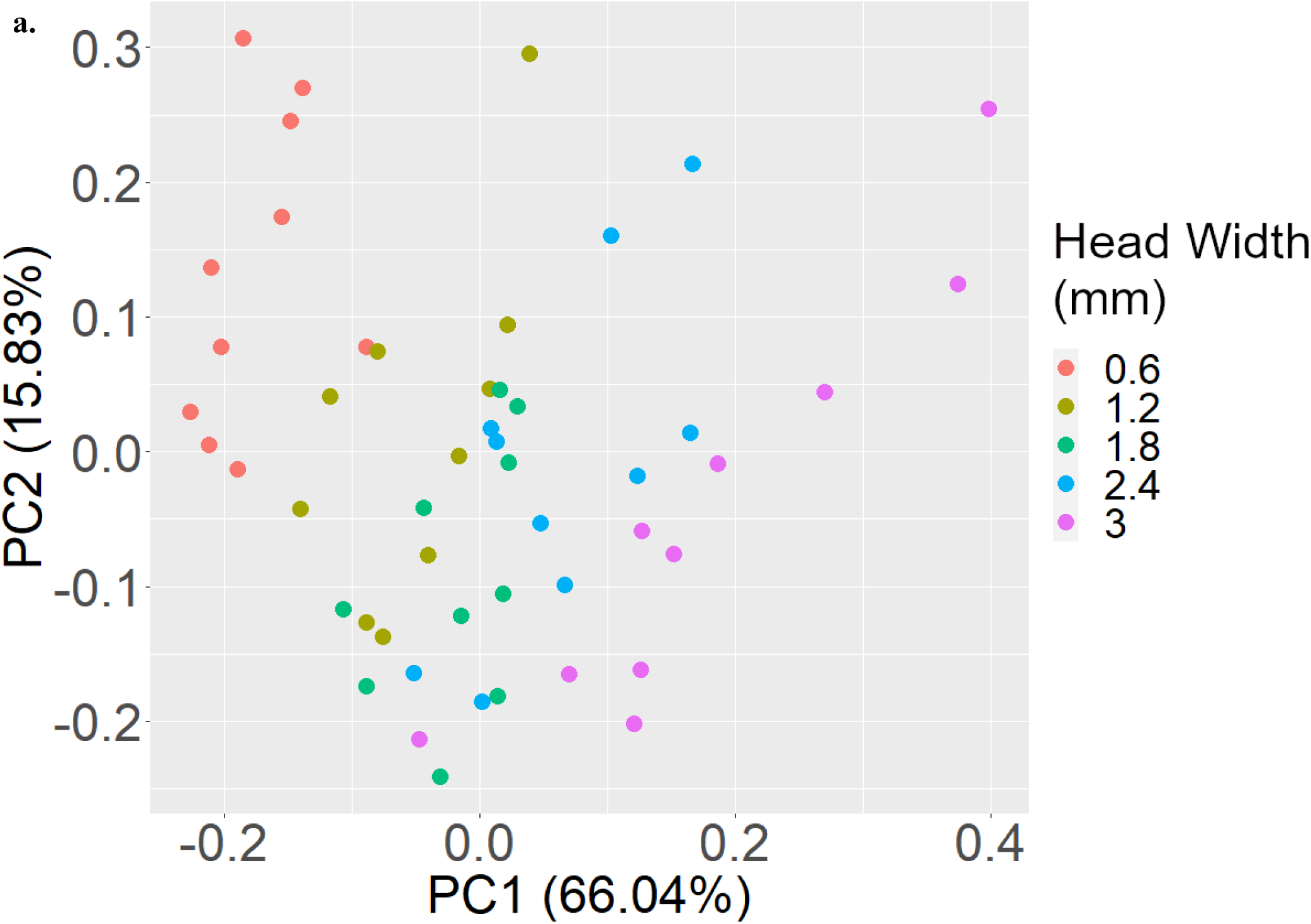
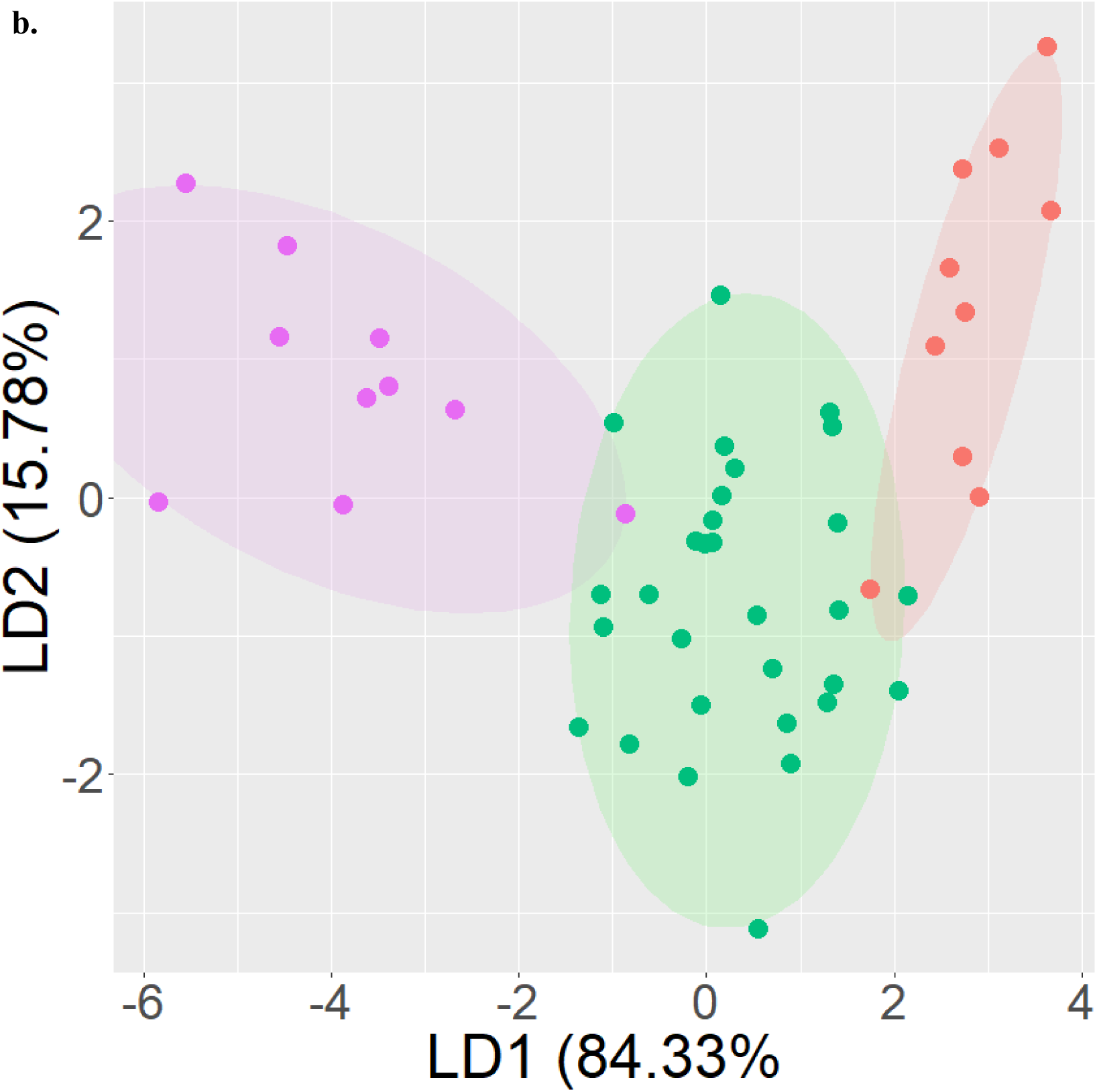
**a.** Principal component analysis plot of all log-transformed compartmental volumes normalized to total brain volume. **b.** Linear discriminant analysis of samples according to worker size group (minims: 0.5-0.7mm, medias: 1.1-2.5mm, majors: 3mm+) based on compartmental volumes (excluding the ROCB, which was colinear with other variables) normalized to total brain size. Classification accuracy = 95.8%. Color indicates worker size group (red = 0.6mm HW, gold = 1.2mm, green = 1.8mm, blue = 2.4mm, purple = 3mm+).

## Discussion

We assessed how estimated sensory and motor requirements for task performance underlying complex agricultural division of labor correlate with quantified variation in brain size and mosaic structure among polymorphic workers of *A. cephalotes*. Behavioral performance challenges in social insects have been typically inferred from interspecific and intraspecific variation in neuropil scaling patterns and general assessments of sensory environments and socioecological influences (Amador-Vargas et al., 2015; Gronenberg, 1999; Gronenberg et al., 1996; Muscedere & Traniello, 2012; O’Donnell, Bulova, Barrett, & Fiocca, 2018), casually correlated with sociobiological characteristics such as colony size, foundation strategy and queen/worker differentiation (for example the reduction in MB volume in solitary compared to social wasps; O’Donnell et al., 2015), and/or diet and life history (Sayol et al., 2020). To the best of our knowledge, our study is the first to computationally estimate the need for brain investment in relation to sensorimotor integration requirements for behavioral performance and establish a correlation with neuroanatomy. The parameters of task performance used to develop our estimate were derived from results of published studies and our own observations rather than subjective or casual assessments of behavior. We evaluated the need for neuropil investment to serve sensory input processing and sensorimotor functions, conservatively evaluated the size-related involvement of polymorphic workers in performing these tasks (Figure 2) and used this metric to generate hypotheses concerning brain evolution. Comparisons of brain compartment volumes indicated significant differences in absolute size and proportional investment across polymorphic task-differentiated workers, supporting the concept that worker neural phenotypes are adaptively designed to meet variation in requirements for task performance. Neuroanatomical scaling patterns broadly correlated with sensory, somatosensory, and integrative information-processing demands associated with the specialized task repertoires of polymorphic workers. Such mosaic scaling is consistent with demands on workers in species with large colonies and worker specialization (Riveros et al., 2012). Notably, our estimate showed a pattern broadly consistent with total MB and MB-P scaling patterns, though the percentage of explained variance was only moderate, lending some support to the idea that selection on brain compartment volume optimizes for task integration requirements though other factors likely influence levels of investment. Such factors may include differences in the amount of tissue required to maintain baseline neural functions across workers that vary in body size and metabolic expenses (Kamhi et al., 2016; Packard, 2020), or other size-related constraints (Finlay & Darlington, 1995; Herculano-Houzel, 2012; Herculano-Houzel et al., 2014; O’Donnell, Bulova, Barrett, & Fiocca, 2018).

### Differentiation of worker neural phenotypes

Principal component and linear discriminant analysis distinguished individual *A. cephalotes* brains from different worker groups on the basis of compartment volumes. LDA demonstrated that the degree to which brains can accurately be classified in terms of worker body size is greater when all media size classes (1.2, 1.8, and 2.4mm) are pooled, but is nevertheless able to distinguish samples belonging to five different groups. These results suggest that medias are readily distinguished from the largest and smallest worker specialists in terms of tissue requirements for task performance but differentiation among size groups of medias is less substantial, and consistent with our estimates (Figure 2).

### Antennal lobes

Medias were predicted to have greater sensory integration needs than other worker groups (Figure 2) due to their large task repertoire and collaterally diverse behavioral challenges such as selecting and harvesting plant material and navigating to and from food sources (Arenas & Roces, 2017; Blanton & Ewel, 1985; Falibene et al., 2015; Groh et al., 2014; Howard, 1987; Howard et al., 1988; Hubbell et al., 1983, 1984; Wilson, 1980a, 1980b). Media workers in the 1.8mm and 2.4mm size groups showed increased AL investment compared with minims and majors, as well as 1.2mm medias. Increased input to the AL from antennal odorant receptors is associated with increased olfactory sensitivity and odor perception (Acebes & Ferrús, 2001; Kuebler et al., 2010). Consistent with their social roles, media worker ALs appear to be enlarged to process more diverse olfactory information, similar to the linkage of AL enlargement and olfaction-based tasks in *Atta vollenweideri* (Kelber et al., 2009; Kleineidam et al., 2005; Kuebler et al., 2010). Nonetheless, little variance in AL volume could be explained by our model, indicating that this method may not be a suitable predictor of investment in primary sensory neuropils, even those sensitive to a wide of array of potential stimuli.

### Mushroom bodies

Medias had the highest relative total MB volume. The trend was consistent for the MB-P and the MB-MC (although differences in proportional MB-MC volume were not significant) but not the MB-LC, which showed significant differences in 1.2 vs. 3mm+ and 1.8 vs. 3mm+ comparisons but no overall trend of increasing in either minims, medias, or majors. The functions of the MB-MCs and the MB-LCs may differ, and interestingly, the medial calyx has been shown to be influenced by experience in bees (Riveros & Gronenberg, 2010). The significant differences we found in total MB volume were thus primarily driven by MB-P volume, an axonal bundle that relays outputs from the MBs to other brain compartments (Gronenberg, 1999), whose volume is more influenced by olfactory experience than visual experience (Heisenberg, 1998). Our results suggest *A. cephalotes* 1.8mm medias require relatively larger MB due to the nature and diversity of their leaf-harvesting task repertoire. These findings are consistent with the broader pattern of task repertoire size and diversity of sensory inputs across polymorphic *A. cephalotes* workers, as factored in our estimate for neural investment demands. In respect to other insect examples of specialist or generalist habits (Farris & Roberts, 2005), the enlargement of the MBs, a crucial sensory processing center, is consistent with the idea that generalists - in this case medias that have a larger task repertoire - require more neural tissue to fulfil their tasks. *A. vollenweideri* workers show decreasing investment in the MB calyces with increasing worker size (Groh et al., 2014). Our data indicate greater MB calyx investment in *A. cephalotes* medias, perhaps due to their generalist plant tissue harvesting, which contrasts with the specialized grass harvesting of *A. vollenweideri*.

### Additional compartmental allometries

We found that CX volume was proportionally largest in minims, and inversely related to worker size. Greater investment in the CX may represent circuitry to enable multisensory navigation within dark three-dimensional labyrinthal fungal comb chambers. Minims mainly perform fungal-gardening tasks that likely rely on non-visual navigational strategies, perhaps involving CX circuitry (Currier et al., 2020; Green et al., 2019; Le Moël et al., 2019; Mamiya et al., 2018; Pisokas et al., 2020; Shiozaki et al., 2020; Sun et al., 2020).

*A. cephalotes* worker OL proportional volume increases with worker size and is highest in majors, which have greater visual acuity (Arganda et al., 2020). The repertoire of *Atta* majors appears limited to defense (Powell & Clark, 2004a; Wilson, 1980a), mainly an extranidal task that likely involves target detection.

### Total brain volume

Total brain volume sharply increased in the largest two worker size groups. When scaled to body size, minims had the largest brains, consistent with Haller’s rule (Rensch, 1956), perhaps due to limits on miniaturization of neural circuits (Beutel et al., 2005; Chittka & Niven, 2009; Groh et al., 2014; Niven & Farris, 2012; Seid et al., 2011). However, Haller’s rule does not apply to hymenopteran species such as some parasitoid wasps smaller in size than *A. cephalotes* minims (Groothuis & Smid, 2017; van der Woude et al., 2013; van der Woude & Smid, 2016). This suggests that *A. cephalotes* fungal gardening and nursing behaviors may demand more neural tissue than that required for parasitoid behavior. Size-adjusted total brain volume showed very little correspondence to estimates of our model of requirements for task performance. Given the more significant correspondence found in some brain compartments and coupling of scaling pattern of compartments to sensory requirements, total brain volume may be a too general and imprecise metric to reflect differences in task performance (Chittka & Niven, 2009; Logan et al., 2018; Muscedere et al., 2014; Muscedere & Traniello, 2012) for *A. cephalotes* workers.

### Conclusions

Our results provide insight into the association of task specializations, their sensorimotor and higher-order processing requirements, and brain evolution. Adaptive brain mosaics in *A. cephalotes* polymorphic workers appear to neuroanatomically support the social organization of agricultural division of labor. We found MB size allometry in media workers that corresponded to estimated sensory processing and sensorimotor demands of leaf harvesting, their principal task. The proportional enlargement of the MBs in this size group supports the hypothesis that task sets having apparently greater requirements for sensory integration, motor coordination, and higher-order processing are associated with significantly greater investment in brain centers underpinning these functions. The relationship of our investment estimate to both MBS and MB-P volumes suggests that our method effectively characterizes neuropils that integrate diverse stimuli and facilitate higher-order processing, but does not adequately explain variances in investment in specialized primary input neuropils. This is likely due to the emphasis our model places on behavioral diversity, and thus the integration and processing of multimodal stimuli, and act frequency.

## Acknowledgements

We gratefully acknowledge the assistance of Dr. Wulfila Gronenberg, Dr. Sam Beshers and Dr. Sean Mullen (manuscript input), Dr. Todd Blute (confocal microscopy), Dr. Christopher Starr and Ricardo Pillai (field work), Andrew Hoadley (colony collection and brain illustrations), and Zach Coto and Frank Azorsa (animal husbandry and manuscript input). This study was supported by National Science Foundation grants IOS 1354291 and IOS 1953393 to JFAT, an award from the Boston University Undergraduate Research Opportunities Program to EMF, and the Department of Biology Brenton R. Lutz Award and the Belamarich Dissertation Writing Award to IBM. Colonies were collected in Trinidad in compliance with the laws of Trinidad and Tobago and imported to USA in compliance with the conditions of permit USDA APHIS P526P-12-04067.

## Author contributions

IBM and JFAT designed the study. IM drafted the manuscript. IM, EMF, and JFAT developed and edited the manuscript. IBM and EMF prepared and imaged brains. EMF measured neuropils and IBM and EMF statistically analyzed volumetric data. JFAT secured funding.

## Notes

### Competing Interest Statement

The authors have declared no competing interest.

## References

Acebes, A., & Ferrús, A. (2001). Increasing the number of synapses modifies olfactory perception in *Drosophila*. The Journal of Neuroscience : The Official Journal of the Society for Neuroscience, 21(16), 6264–6273. PubMed. https://doi.org/10.1523/JNEUROSCI.21-16-06264.2001

Adolphs, R. (2003). Cognitive neuroscience of human social behaviour. Nature Reviews Neuroscience, 4(3), 165–178. https://doi.org/10.1038/nrn1056

Aiello, L. C., & Wheeler, P. (1995). The expensive-tissue hypothesis: The brain and the digestive system in human and primate evolution. Current Anthropology, 36(2), 199–221. https://doi.org/10.1086/204350

Amador-Vargas, S., Gronenberg, W., Wcislo, W. T., & Mueller, U. (2015). Specialization and group size: Brain and behavioural correlates of colony size in ants lacking morphological castes. Proceedings of the Royal Society B: Biological Sciences, 282(1801), 20142502. https://doi.org/10.1098/rspb.2014.2502

Arenas, A., & Roces, F. (2016a). Gardeners and midden workers in leaf-cutting ants learn to avoid plants unsuitable for the fungus at their worksites. Animal Behaviour, 115, 167–174. https://doi.org/10.1016/j.anbehav.2016.03.016

Arenas, A., & Roces, F. (2016b). Learning through the waste: Olfactory cues from the colony refuse influence plant preferences in foraging leaf-cutting ants. The Journal of Experimental Biology, 219(16), 2490–2496. https://doi.org/10.1242/jeb.139568

Arenas, A., & Roces, F. (2017). Avoidance of plants unsuitable for the symbiotic fungus in leaf-cutting ants: Learning can take place entirely at the colony dump. PLOS ONE, 12(3), e0171388. https://doi.org/10.1371/journal.pone.0171388

Arganda, S., Hoadley, A. P., Razdan, E. S., Muratore, I. B., & Traniello, J. F. A. (2020). The neuroplasticity of division of labor: Worker polymorphism, compound eye structure and brain organization in the leafcutter ant *Atta cephalotes*. Journal of Comparative Physiology A. https://doi.org/10.1101/2020.03.04.975110

Armitage, S. A. O., Fernández-Marín, H., Boomsma, J. J., & Wcislo, W. T. (2016). Slowing them down will make them lose: A role for attine ant crop fungus in defending pupae against infections? The Journal of Animal Ecology, 85(5), 1210–1221. https://doi.org/10.1111/1365-2656.12543

Banks, A. N., & Srygley, R. B. (2003). Orientation by magnetic field in leaf-cutter ants, *Atta colombica* (Hymenoptera: Formicidae). Ethology, 109(10), 835–846. https://doi.org/10.1046/j.0179-1613.2003.00927.x

Bass, M., & Cherrett, J. M. (1996). Leaf-cutting ants (formicidae, attini) prune their fungus to increase and direct its productivity. Functional Ecology, 10(1), 55–61. https://doi.org/10.2307/2390262

Beshers, S. N., & Fewell, J. H. (2001). Models of division of labor in social insects. Annual Review of Entomology, 46(1), 413–440. https://doi.org/10.1146/annurev.ento.46.1.413

Beutel, R. G., Pohl, H., & Hünefeld, F. (2005). Strepsipteran brains and effects of miniaturization (Insecta). Arthropod Structure & Development, 34(3), 301–313. https://doi.org/10.1016/j.asd.2005.03.001

Blanton, C. M., & Ewel, J. J. (1985). Leaf-cutting ant herbivory in successional and agricultural tropical ecosystems. Ecology, 66(3), 861–869. https://doi.org/10.2307/1940548

Bollazzi, M., & Roces, F. (2002). Thermal preference for fungus culturing and brood location by workers of the thatching grass-cutting ant *Acromyrmex heyeri*. Insectes Sociaux, 49(2), 153–157. https://doi.org/10.1007/s00040-002-8295-x

Boogert, N. J., Madden, J. R., Morand-Ferron, J., & Thornton, A. (2018). Measuring and understanding individual differences in cognition. Philosophical Transactions of the Royal Society B: Biological Sciences, 373(1756), 20170280. https://doi.org/10.1098/rstb.2017.0280

Camargo, R. da S. (2015). Foraging behavior of leaf cutting ants: How do workers search for their food? Sociobiology, 62(3), 347–350. https://doi.org/10.13102/sociobiology.v62i3.714

Chittka, L., & Niven, J. (2009). Are bigger brains better? Current Biology, 19(21), R995–R1008. https://doi.org/10.1016/j.cub.2009.08.023

Currie, C. R., & Stuart, A. E. (2001). Weeding and grooming of pathogens in agriculture by ants. Proceedings of the Royal Society of London. Series B: Biological Sciences, 268(1471), 1033–1039. https://doi.org/10.1098/rspb.2001.1605

Currier, T. A., Matheson, A. M., & Nagel, K. I. (2020). Encoding and control of orientation to airflow by a set of *Drosophila* fan-shaped body neurons. ELife, 9, e61510. https://doi.org/10.7554/eLife.61510

DeCasien, A. R., & Higham, J. P. (2019). Primate mosaic brain evolution reflects selection on sensory and cognitive specialization. Nature Ecology & Evolution, 3(10), 1483–1493. https://doi.org/10.1038/s41559-019-0969-0

DeCasien, A. R., Williams, S. A., & Higham, J. P. (2017). Primate brain size is predicted by diet but not sociality. Nature Ecology & Evolution, 1(5). https://doi.org/10.1038/s41559-017-0112

Dunbar, R. I. M. (1998). The social brain hypothesis. Evolutionary Anthropology: Issues, News, and Reviews, 6(5), 178–190. https://doi.org/10.1002/(sici)1520-6505(1998)6:5<178::aid-evan5>3.0.co;2-8

Dunbar, R. I. M. (2009). The social brain hypothesis and its implications for social evolution. Annals of Human Biology, 36(5), 562–572. https://doi.org/10.1080/03014460902960289

Dupuis, E. C., & Harrison, J. F. (2017). Trunk trail maintenance in leafcutter ants: Caste involvement and effects of obstacle type and size on path clearing in *Atta cephalotes*. Insectes Sociaux, 64(2), 189–196. https://doi.org/10.1007/s00040-016-0530-y

Durst, C., Eichmüller, S., & Menzel, R. (1994). Development and experience lead to increased volume of subcompartments of the honeybee mushroom body. Behavioral and Neural Biology, 62(3), 259–263. https://doi.org/10.1016/s0163-1047(05)80025-1

Evison, S. E. F., Hart, A. G., & Jackson, D. E. (2008). Minor workers have a major role in the maintenance of leafcutter ant pheromone trails. Animal Behaviour, 75(3), 963–969. https://doi.org/10.1016/j.anbehav.2007.07.013

Fahrbach, S. E. (2006). Structure of the mushroom bodies of the insect brain. Annual Review of Entomology, 51(1), 209–232. https://doi.org/10.1146/annurev.ento.51.110104.150954

Falibene, A., Roces, F., & Rössler, W. (2015). Long-term avoidance memory formation is associated with a transient increase in mushroom body synaptic complexes in leaf-cutting ants. Frontiers in Behavioral Neuroscience, 9, 84–84. PubMed. https://doi.org/10.3389/fnbeh.2015.00084

Farias, A. P., Camargo, R. da S., Andrade Sousa, K. K., Caldato, N., & Forti, L. C. (2020). Nest architecture and colony growth of *Atta bisphaerica* grass-cutting ants. Insects, 11(11), 741. https://doi.org/10.3390/insects11110741

Farris, S. M. (2011). Are mushroom bodies cerebellum-like structures? Arthropod Structure & Development, 40(4), 368–379. https://doi.org/10.1016/j.asd.2011.02.004

Farris, S. M., & Roberts, N. S. (2005). Coevolution of generalist feeding ecologies and gyrencephalic mushroom bodies in insects. Proceedings of the National Academy of Sciences, 102(48), 17394–17399. https://doi.org/10.1073/pnas.0508430102

Feener, D. H., & Moss, K. A. G. (1990). Defense against parasites by hitchhikers in leaf-cutting ants: A quantitative assessment. Behavioral Ecology and Sociobiology, 26(1), 17–29. https://doi.org/10.1007/bf00174021

Feinerman, O., & Traniello, J. F. A. (2015). Social complexity, diet, and brain evolution: Modeling the effects of colony size, worker size, brain size, and foraging behavior on colony fitness in ants. Behavioral Ecology and Sociobiology, 70(7), 1063–1074. https://doi.org/10.1007/s00265-015-2035-5

Finlay, B., & Darlington, R. (1995). Linked regularities in the development and evolution of mammalian brains. Science, 268(5217), 1578–1584. https://doi.org/10.1126/science.7777856

Franks, N. R., & Sendova-Franks, A. B. (1992). Brood sorting by ants: Distributing the workload over the work-surface. Behavioral Ecology and Sociobiology, 30(2), 109–123. https://doi.org/10.1007/BF00173947

Godfrey, R. K., & Gronenberg, W. (2019). Brain evolution in social insects: Advocating for the comparative approach. Journal of Comparative Physiology A, 205(1), 13–32. https://doi.org/10.1007/s00359-019-01315-7

Goes, A. C., Barcoto, M. O., Kooij, P. W., Bueno, O. C., & Rodrigues, A. (2020). How do leaf-cutting ants recognize antagonistic microbes in their fungal crops? Frontiers in Ecology and Evolution, 8. https://doi.org/10.3389/fevo.2020.00095

Gordon, D. G., Ilieş, I., & Traniello, J. F. A. (2017). Behavior, brain, and morphology in a complex insect society: Trait integration and social evolution in the exceptionally polymorphic ant *Pheidole rhea*. Behavioral Ecology and Sociobiology, 71(11), 166. https://doi.org/10.1007/s00265-017-2396-z

Green, J., Vijayan, V., Mussells Pires, P., Adachi, A., & Maimon, G. (2019). A neural heading estimate is compared with an internal goal to guide oriented navigation. Nature Neuroscience, 22(9), 1460–1468. https://doi.org/10.1038/s41593-019-0444-x

Green, P. W. C., & Kooij, P. W. (2018). The role of chemical signalling in maintenance of the fungus garden by leaf-cutting ants. Chemoecology, 28(3), 101–107. https://doi.org/10.1007/s00049-018-0260-x

Groh, C., Kelber, C., Grübel, K., & Rössler, W. (2014). Density of mushroom body synaptic complexes limits intraspecies brain miniaturization in highly polymorphic leaf-cutting ant workers. Proceedings of the Royal Society B: Biological Sciences, 281(1785), 20140432–20140432. PubMed. https://doi.org/10.1098/rspb.2014.0432

Gronenberg, W. (1999). Modality-specific segregation of input to ant mushroom bodies. Brain, Behavior and Evolution, 54(2), 85–95. https://doi.org/10.1159/000006615

Gronenberg, W. (2001). Subdivisions of hymenopteran mushroom body calyces by their afferent supply. Journal of Comparative Neurology, 435(4), 474–489. https://doi.org/10.1002/cne.1045

Gronenberg, W., Heeren, S., & Hölldobler, B. (1996). Age-dependent and task-related morphological changes in the brain and the mushroom bodies of the ant *Camponotus floridanus*. Journal of Experimental Biology, 199(9), 2011–2019.

Gronenberg, W., & Riveros, A. J. (2009). Social brains and behavior: Past and present. Organization of Insect Societies From Genome to Sociocomplexity.

Groothuis, J., & Smid, H. M. (2017). Nasonia Parasitic Wasps Escape from Haller’s Rule by Diphasic, Partially Isometric Brain-Body Size Scaling and Selective Neuropil Adaptations. Brain, Behavior and Evolution, 90(3), 243–254. https://doi.org/10.1159/000480421

Hart, A. G., & Ratnieks, F. L. W. (2001). Task partitioning, division of labour and nest compartmentalisation collectively isolate hazardous waste in the leafcutting ant *Atta cephalotes*. Behavioral Ecology and Sociobiology, 49(5), 387–392. https://doi.org/10.1007/s002650000312

Heisenberg, M. (1998). What do the mushroom bodies do for the insect brain? An introduction. Learning & Memory, 5(1), 1–10. https://doi.org/10.1101/lm.8.1.1

Herculano-Houzel, S. (2012). The remarkable, yet not extraordinary, human brain as a scaled-up primate brain and its associated cost. Proceedings of the National Academy of Sciences, 109(Supplement 1), 10661–10668.

Herculano-Houzel, S., Manger, P. R., & Kaas, J. H. (2014). Brain scaling in mammalian evolution as a consequence of concerted and mosaic changes in numbers of neurons and average neuronal cell size. Frontiers in Neuroanatomy, 8. https://doi.org/10.3389/fnana.2014.00077

Hölldobler, B., & Wilson, E. O. (1990). The Ants. Harvard University Press.

Hölldobler, B., & Wilson, E. O. (2010). The leafcutter ants: Civilization by instinct. W. W. Norton & Company.

Howard, J. J. (1987). Leafcutting ant diet selection: The role of nutrients, water, and secondary chemistry. Ecology, 68(3), 503–515. https://doi.org/10.2307/1938455

Howard, J. J., Cazin, J., & Wiemer, D. F. (1988). Toxicity of terpenoid deterrents to the leafcutting ant *Atta cephalotes* and its mutualistic fungus. Journal of Chemical Ecology, 14(1), 59–69. https://doi.org/10.1007/bf01022531

Howard, J. J., Henneman, L. M., Cronin, G., Fox, J. A., & Hormiga, G. (1996). Conditioning of scouts and recruits during foraging by a leaf-cutting ant, *Atta colombica*. Animal Behaviour, 52(2), 299–306. https://doi.org/10.1006/anbe.1996.0175

Hubbell, S. P., Howard, J. J., & Wiemer, D. F. (1984). Chemical leaf repellency to an attine ant: Seasonal distribution among potential host plant species. Ecology, 65(4), 1067–1076. https://doi.org/10.2307/1938314

Hubbell, S. P., Wiemer, D. F., & Adejare, A. (1983). An antifungal terpenoid defends a neotropical tree (Hymenaea) against attack by fungus-growing ants (*Atta*). Oecologia, 60(3), 321–327. https://doi.org/10.1007/bf00376846

Ilieş, I., Muscedere, M. L., & Traniello, J. F. A. (2015). Neuroanatomical and morphological trait clusters in the ant genus *Pheidole*: Evidence for modularity and integration in brain structure. Brain, Behavior and Evolution, 85(1), 63–76. https://doi.org/10.1159/000370100

Jaffe, K., & Perez, E. (1989). Comparative study of brain morphology in ants. Brain, Behavior and Evolution, 33(1), 25–33. https://doi.org/10.1159/000115895

Kamhi, J. F., Barron, A. B., & Narendra, A. (2020). Vertical lobes of the mushroom bodies are essential for view-based navigation in australian *Myrmecia* ants. Current Biology, 30(17), 3432–3437.e3. https://doi.org/10.1016/j.cub.2020.06.030

Kamhi, J. F., Gronenberg, W., Robson, S. K. A., & Traniello, J. F. A. (2016). Social complexity influences brain investment and neural operation costs in ants. Proceedings of the Royal Society B: Biological Sciences, 283(1841), 20161949. PubMed. https://doi.org/10.1098/rspb.2016.1949

Kamhi, J. F., Ilieş, I., & Traniello, J. F. A. (2019). Social complexity and brain evolution: Comparative analysis of modularity and integration in ant brain organization. *Brain*, Behavior and Evolution, 93(1), 4–18. https://doi.org/10.1159/000497267

Kelber, C., Rössler, W., & Kleineidam, C. J. (2009). Phenotypic plasticity in number of glomeruli and sensory innervation of the antennal lobe in leaf-cutting ant workers (*A. vollenweideri*). Developmental Neurobiology, 70(4), 222–234. https://doi.org/10.1002/dneu.20782

Khadempour, L. (2018). Microbial mediation of herbivory in leaf-cutter ant fungus gardens [Ph.D.]. The University of Wisconsin - Madison.

Khadempour, L., Kyle, J. E., Webb-Robertson, B.-J. M., Nicora, C. D., Smith, F. B., Smith, R. D., Lipton, M. S., Currie, C. R., Baker, E. S., & Burnum-Johnson, K. E. (2020). From plants to ants: Fungal modification of leaf lipids for nutrition and communication in the leaf-cutter ant fungal garden ecosystem. BioRxiv, 2020.07.28.224139. https://doi.org/10.1101/2020.07.28.224139

Khalife, A., Keller, R. A., Billen, J., Hita Garcia, F., Economo, E. P., & Peeters, C. (2018). Skeletomuscular adaptations of head and legs of *Melissotarsus* ants for tunnelling through living wood. Frontiers in Zoology, 15(1), 30. https://doi.org/10.1186/s12983-018-0277-6

Kleineidam, C. J., Obermayer, M., Halbich, W., & Rössler, W. (2005). A macroglomerulus in the antennal lobe of leaf-cutting ant workers and its possible functional significance. Chemical Senses, 30(5), 383–392. https://doi.org/10.1093/chemse/bji033

Kooij, P. W., Pullens, J. W. M., Boomsma, J. J., & Schiøtt, M. (2016). Ant mediated redistribution of a xyloglucanase enzyme in fungus gardens of *Acromyrmex echinatior*. BMC Microbiology, 16(1), 81. https://doi.org/10.1186/s12866-016-0697-4

Kuebler, L. S., Kelber, C., & Kleineidam, C. J. (2010). Distinct antennal lobe phenotypes in the leaf-cutting ant (*Atta vollenweideri*). The Journal of Comparative Neurology, 518(3), 352–365. https://doi.org/10.1002/cne.22217

Kwaku, K. M., Gonick, E. A., Ostapovich, E. M., & Weinberg, I. P. (2020). The frequency of leaf transfer in *Atta cephalotes* along horizontal and vertical surfaces near the bases of trees. Insectes Sociaux, 67(4), 481–486. https://doi.org/10.1007/s00040-020-00784-3

Le Moël, F., Stone, T., Lihoreau, M., Wystrach, A., & Webb, B. (2019). The central complex as a potential substrate for vector based navigation. Frontiers in Psychology, 10. https://doi.org/10.3389/fpsyg.2019.00690

Lenoir, A., D’Ettorre, P., Errard, C., & Hefetz, A. (2001). Chemical ecology and social parasitism in ants. Annual Review of Entomology, 46(1), 573–599. https://doi.org/10.1146/annurev.ento.46.1.573

Liaw, A., & Wiener, M. (2001). Classification and regression by randomforest. Forest, 23.

Lihoreau, M., Dubois, T., Gomez-Moracho, T., Kraus, S., Monchanin, C., & Pasquaretta, C. (2019). Putting the ecology back into insect cognition research. Advances in Insect Physiology, 1–25. https://doi.org/10.1016/bs.aiip.2019.08.002

Lihoreau, M., Latty, T., & Chittka, L. (2012). An exploration of the social brain hypothesis in insects. Frontiers in Physiology, 3, 442–442. PubMed. https://doi.org/10.3389/fphys.2012.00442

Logan, C., Avin, S., Boogert, N., Buskell, A., Cross, F. R., Currie, A., Jelbert, S., Lukas, D., Mares, R., Navarrete, A. F., Shigeno, S., & Montgomery, S. (2018). Beyond brain size: Uncovering the neural correlates of behavioral and cognitive specialization. https://doi.org/10.17863/CAM.25916

Markl, H. (1965). Stridulation in leaf-cutting ants. Science (New York, N.Y.), 149(3690), 1392–1393. https://doi.org/10.1126/science.149.3690.1392

Moll, K., Roces, F., & Federle, W. (2013). How load-carrying ants avoid falling over: Mechanical stability during foraging in *Atta vollenweideri* grass-cutting ants. PloS One, 8(1), e52816. https://doi.org/10.1371/journal.pone.0052816

Mueller, U. G. (2002). Ant versus fungus versus mutualism: Ant-cultivar conflict and the deconstruction of the attine ant-fungus symbiosis. The American Naturalist, 160 Suppl 4, S67–98. https://doi.org/10.1086/342084

Muratore, I. B., Ilieş, I., Huzar, A. K., Zaidi, F. H., & Traniello, J. F. A. (in prep.). Morphological scaling and division of labor in a socially complex agricultural ant.

Muratore, I. B., & Traniello, J. F. A. (2020). Fungus-growing ants: Models for the integrative analysis of cognition and brain evolution. Frontiers in Behavioral Neuroscience, 14, 599234–599234. PubMed. https://doi.org/10.3389/fnbeh.2020.599234

Muscedere, M. L., Gronenberg, W., Moreau, C. S., & Traniello, J. F. A. (2014). Investment in higher order central processing regions is not constrained by brain size in social insects. Proceedings of the Royal Society B: Biological Sciences, 281(1784), 20140217. https://doi.org/10.1098/rspb.2014.0217

Muscedere, M. L., & Traniello, J. F. A. (2012). Division of labor in the hyperdiverse ant genus *Pheidole* is associated with distinct subcaste- and age-related patterns of worker brain organization. PloS One, 7(2), e31618–e31618. PubMed. https://doi.org/10.1371/journal.pone.0031618

Navarrete, A. F., Reader, S. M., Street, S. E., Whalen, A., & Laland, K. N. (2016). The coevolution of innovation and technical intelligence in primates. Philosophical Transactions of the Royal Society of London. Series B, Biological Sciences, 371(1690), 20150186. PubMed. https://doi.org/10.1098/rstb.2015.0186

Nichols-Orians, C. M., & Schultz, J. C. (1989). Leaf toughness affects leaf harvesting by the leaf cutter ant, *Atta cephalotes* (l.) (Hymenoptera: Formicidae). Biotropica, 21(1), 80–83. https://doi.org/10.2307/2388446

Niven, J. E., & Farris, S. M. (2012). Miniaturization of nervous systems and neurons. Current Biology, 22(9), R323–R329. https://doi.org/10.1016/j.cub.2012.04.002

Norman, V. C., Butterfield, T., Drijfhout, F., Tasman, K., & Hughes, W. O. H. (2017). Alarm pheromone composition and behavioral activity in fungus-growing ants. Journal of Chemical Ecology, 43(3), 225–235. https://doi.org/10.1007/s10886-017-0821-4

O’Donnell, S., Bulova, S., Barrett, M., & von Beeren, C. (2018). Brain investment under colony-level selection: Soldier specialization in *Eciton* army ants (Formicidae: Dorylinae). BMC Zoology, 3(1), 3. https://doi.org/10.1186/s40850-018-0028-3

O’Donnell, S., Bulova, S. J., Barrett, M., & Fiocca, K. (2018). Size constraints and sensory adaptations affect mosaic brain evolution in paper wasps (Vespidae: Epiponini). Biological Journal of the Linnean Society, 123(2), 302–310. https://doi.org/10.1093/biolinnean/blx150

O’Donnell, S., Bulova, S. J., DeLeon, S., Khodak, P., Miller, S., & Sulger, E. (2015). Distributed cognition and social brains: Reductions in mushroom body investment accompanied the origins of sociality in wasps (Hymenoptera: Vespidae). Proceedings of the Royal Society B: Biological Sciences, 282(1810), 20150791. PubMed. https://doi.org/10.1098/rspb.2015.0791

Orr, M. R. (1992). Parasitic flies (Diptera: Phoridae) influence foraging rhythms and caste division of labor in the leaf-cutter ant, *Atta cephalotes* (Hymenoptera: Formicidae). Behavioral Ecology and Sociobiology, 30(6), 395–402. https://doi.org/10.1007/BF00176174

Packard, G. C. (2020). Rethinking the metabolic allometry of ants. Evolutionary Ecology, 34(2), 149–161. https://doi.org/10.1007/s10682-020-10033-5

Pisokas, I., Heinze, S., & Webb, B. (2020). The head direction circuit of two insect species. ELife, 9, e53985. PubMed. https://doi.org/10.7554/eLife.53985

Poulsen, M., Bot, A. N., Nielsen, M. G., & Boomsma, J. J. (2002). Experimental evidence for the costs and hygienic significance of the antibiotic metapleural gland secretion in leaf-cutting ants. Behavioral Ecology and Sociobiology, 52(2), 151–157. https://doi.org/10.1007/s00265-002-0489-8

Powell, S., & Clark, E. (2004a). Combat between large derived societies: A subterranean army ant established as a predator of mature leaf-cutting ant colonies. Insectes Sociaux, 4(51), 342–351. https://doi.org/10.1007/s00040-004-0752-2

Powell, S., & Clark, E. (2004b). Combat between large derived societies: A subterranean army ant established as a predator of mature leaf-cutting ant colonies. Insectes Sociaux, 51(4), 342–351. https://doi.org/10.1007/s00040-004-0752-2

Rensch, B. (1956). Increase of learning capability with increase of brain-size. The American Naturalist, 90(851), 81–95. https://doi.org/10.1086/281911

Reséndiz-Benhumea, G. M., Sangati, E., Sangati, F., Keshmiri, S., & Froese, T. (2021). Shrunken Social Brains? A Minimal Model of the Role of Social Interaction in Neural Complexity. Frontiers in Neurorobotics, 15. https://doi.org/10.3389/fnbot.2021.634085

Riveros, A. J., & Gronenberg, W. (2010). Brain allometry and neural plasticity in the bumblebee *Bombus occidentalis*. Brain, Behavior and Evolution, 75(2), 138–148. PubMed. https://doi.org/10.1159/000306506

Riveros, A. J., Seid, M. A., & Wcislo, W. T. (2012). Evolution of brain size in class-based societies of fungus-growing ants (Attini). Animal Behaviour, 83(4), 1043–1049. https://doi.org/10.1016/j.anbehav.2012.01.032

Roces, F., & Núñez, JosuéA. (1995). Thermal sensitivity during brood care in workers of two *Camponotus* ant species: Circadian variation and its ecological correlates. Journal of Insect Physiology, 41(8), 659–669. https://doi.org/10.1016/0022-1910(95)00019-Q

Roces, F., Tautz, J., & Hölldobler, B. (1993). Stridulation in leaf-cutting ants. Naturwissenschaften, 80(11), 521–524. https://doi.org/10.1007/BF01140810

Rodrigues, A., Bacci Jr, M., Mueller, U. G., Ortiz, A., & Pagnocca, F. C. (2008). Microfungal “weeds” in the leafcutter ant symbiosis. Microbial Ecology, 56(4), 604–614. https://doi.org/10.1007/s00248-008-9380-0

Römer, D., Bollazzi, M., & Roces, F. (2017). Carbon dioxide sensing in an obligate insect-fungus symbiosis: CO2 preferences of leaf-cutting ants to rear their mutualistic fungus. PLOS ONE, 12(4), e0174597. https://doi.org/10.1371/journal.pone.0174597

Römer, D., Bollazzi, M., & Roces, F. (2018). Carbon dioxide sensing in the social context: Leaf-cutting ants prefer elevated CO2 levels to tend their brood. Journal of Insect Physiology, 108, 40–47. https://doi.org/10.1016/j.jinsphys.2018.05.007

Römer, D., & Roces, F. (2014). Nest enlargement in leaf-cutting ants: Relocated brood and fungus trigger the excavation of new chambers. PloS One, 9(5), e97872. https://doi.org/10.1371/journal.pone.0097872

Rytter, W., & Shik, J. Z. (2016). Liquid foraging behaviour in leafcutting ants: The lunchbox hypothesis. Animal Behaviour, 117, 179–186. https://doi.org/10.1016/j.anbehav.2016.04.022

Saverschek, N., & Roces, F. (2011). Foraging leafcutter ants: Olfactory memory underlies delayed avoidance of plants unsuitable for the symbiotic fungus. Animal Behaviour, 82(3), 453–458. https://doi.org/10.1016/j.anbehav.2011.05.015

Sayol, F., Collado, M. Á., Garcia-Porta, J., Seid, M. A., Gibbs, J., Agorreta, A., San Mauro, D., Raemakers, I., Sol, D., & Bartomeus, I. (2020). Feeding specialization and longer generation time are associated with relatively larger brains in bees. Proceedings of the Royal Society B: Biological Sciences, 287(1935), 20200762–20200762. PubMed. https://doi.org/10.1098/rspb.2020.0762

Schultner, E., & Pulliainen, U. (2020). Brood recognition and discrimination in ants. Insectes Sociaux, 67(1), 11–34. https://doi.org/10.1007/s00040-019-00747-3

Segre, P. S., & Taylor, E. D. (2019). Large ants do not carry their fair share: Maximal load-carrying performance of leaf-cutter ants (*Atta cephalotes*). Journal of Experimental Biology, 222(12). https://doi.org/10.1242/jeb.199240

Seid, M. A., Castillo, A., & Wcislo, W. T. (2011). The allometry of brain miniaturization in ants. Brain, Behavior and Evolution, 77(1), 5–13. https://doi.org/10.1159/000322530

Shiozaki, H. M., Ohta, K., & Kazama, H. (2020). A Multi-regional Network Encoding Heading and Steering Maneuvers in Drosophila. Neuron, 106(1), 126–141.e5. https://doi.org/10.1016/j.neuron.2020.01.009

Simons, M., & Tibbetts, E. (2019). Insects as models for studying the evolution of animal cognition. Current Opinion in Insect Science, 34, 117–122. https://doi.org/10.1016/j.cois.2019.05.009

Smaers, J. B., & Soligo, C. (2013). Brain reorganization, not relative brain size, primarily characterizes anthropoid brain evolution. Proceedings of the Royal Society B: Biological Sciences, 280(1759), 20130269. https://doi.org/10.1098/rspb.2013.0269

Strausfeld, N. J. (2012). Arthropod Brains.

Sulger, E., McAloon, N., Bulova, S. J., Sapp, J., & O’Donnell, S. (2014). Evidence for adaptive brain tissue reduction in obligate social parasites (Polyergus mexicanus) relative to their hosts (Formica fusca). Biological Journal of the Linnean Society, 113(2), 415–422. https://doi.org/10.1111/bij.12375

Sun, X., Yue, S., & Mangan, M. (2020). Author response: A decentralised neural model explaining optimal integration of navigational strategies in insects. https://doi.org/10.7554/elife.54026.sa2

Thiele, T., Kost, C., Roces, F., & Wirth, R. (2014). Foraging leaf-cutting ants learn to reject *Vitis vinifera ssp. Vinifera* plants that emit herbivore-induced volatiles. Journal of Chemical Ecology, 40(6), 617–620. https://doi.org/10.1007/s10886-014-0460-y

Tragust, S., Ugelvig, L. V., Chapuisat, M., Heinze, J., & Cremer, S. (2013). Pupal cocoons affect sanitary brood care and limit fungal infections in ant colonies. BMC Evolutionary Biology, 13, 225. https://doi.org/10.1186/1471-2148-13-225

van der Woude, E., & Smid, H. M. (2016). How to escape from haller’s rule: Olfactory system complexity in small and large Trichogramma evanescens parasitic wasps. Journal of Comparative Neurology, 524(9), 1876–1891. https://doi.org/10.1002/cne.23927

van der Woude, E., Smid, H. M., Chittka, L., & Huigens, M. E. (2013). Breaking Haller’s Rule: Brain-Body Size Isometry in a Minute Parasitic Wasp. Brain, Behavior and Evolution, 81(2), 86–92. https://doi.org/10.1159/000345945

Wartel, A., Lindenfors, P., & Lind, J. (2019). Whatever you want: Inconsistent results are the rule, not the exception, in the study of primate brain evolution. PLOS ONE, 14(7), e0218655. https://doi.org/10.1371/journal.pone.0218655

Wehner, R., Fukushi, T., & Isler, K. (2007). On being small: Brain allometry in ants. Brain, Behavior and Evolution, 69(3), 220–228. https://doi.org/10.1159/000097057

Wilson, E. O. (1980a). Caste and division of labor in leaf-cutter ants (Hymenoptera, Formicidae, *Atta*). 1. The overall pattern in *Atta-Sexdens*. Behavioral Ecology and Sociobiology, 7(2), 143–156. https://doi.org/10.1007/BF00299520

Wilson, E. O. (1980b). Caste and division of labor in leaf-cutter ants (Hymenoptera: Formicidae: *Atta*) II. The ergonomic optimization of leaf cutting. Behavioral Ecology and Sociobiology, 7(2), 157–165. https://doi.org/10.1007/bf00299521

Wilson, E. O. (1983). Caste and division of labor in leaf-cutter ants (Hymenoptera: Formicidae: Atta). Behavioral Ecology and Sociobiology, 14(1), 55–60. https://doi.org/10.1007/BF00366656

