## Supplementary Materials for "Behavioral performance requirements for division of labor influence adaptive brain mosaicism in a socially complex ant"

**Supplementary Table 1.** Absolute volumes (µm^3^) of brain compartments and sums. Bin indicates HW(mm) category, divided into three groups (bins) for media workers. HW indicates actual size measurements of sampled individuals according to bin grouping.

| Colony | ID | Bin | SUM | OL | AL | MB-MC | MB-LC | MB-P | CB | SEG | ROCB |
| --- | --- | --- | --- | --- | --- | --- | --- | --- | --- | --- | --- |
| Ac20 | 006 | 0.6 | 1.115E+05 | 1.423E+03 | 6.226E+03 | 8.743E+03 | 9.974E+03 | 1.239E+04 | 1.852E+03 | 1.688E+04 | 5.402E+04 |
| Ac20 | 056 | 1.2 | 1.717E+05 | 3.467E+03 | 1.187E+04 | 1.749E+04 | 1.739E+04 | 2.272E+04 | 2.257E+03 | 3.338E+04 | 6.317E+04 |
| Ac20 | 055 | 1.2 | 2.103E+05 | 3.034E+03 | 1.415E+04 | 2.071E+04 | 2.205E+04 | 2.671E+04 | 3.007E+03 | 4.269E+04 | 7.795E+04 |
| Ac20 | 036 | 1.8 | 2.207E+05 | 5.413E+03 | 1.699E+04 | 2.268E+04 | 2.181E+04 | 3.144E+04 | 1.960E+03 | 3.262E+04 | 8.780E+04 |
| Ac20 | 34 | 1.8 | 2.391E+05 | 5.452E+03 | 1.863E+04 | 2.315E+04 | 2.331E+04 | 2.220E+04 | 1.983E+03 | 5.049E+04 | 9.390E+04 |
| Ac20 | 033 | 1.8 | 2.524E+05 | 4.961E+03 | 2.212E+04 | 2.646E+04 | 2.662E+04 | 2.615E+04 | 1.908E+03 | 4.082E+04 | 1.034E+05 |
| Ac20 | 29 | 1.8 | 2.727E+05 | 5.908E+03 | 1.676E+04 | 2.844E+04 | 2.717E+04 | 3.164E+04 | 2.003E+03 | 4.206E+04 | 1.187E+05 |
| Ac21 | 36 | 1.8 | 2.753E+05 | 7.859E+03 | 1.883E+04 | 2.672E+04 | 2.790E+04 | 3.169E+04 | 3.721E+03 | 4.760E+04 | 1.110E+05 |
| Ac21 | 30 | 1.8 | 3.027E+05 | 6.364E+03 | 1.938E+04 | 3.023E+04 | 2.972E+04 | 4.006E+04 | 4.393E+03 | 5.371E+04 | 1.188E+05 |
| Ac20 | 028 | 3 | 3.078E+05 | 1.227E+04 | 2.385E+04 | 3.014E+04 | 3.127E+04 | 2.969E+04 | 2.251E+03 | 5.725E+04 | 1.211E+05 |
| Ac20 | 21 | 2.4 | 3.404E+05 | 1.063E+04 | 3.295E+04 | 3.159E+04 | 3.481E+04 | 3.704E+04 | 1.626E+03 | 5.987E+04 | 1.318E+05 |
| Ac21 | 28 | 2.4 | 3.563E+05 | 9.640E+03 | 3.240E+04 | 3.314E+04 | 3.460E+04 | 4.202E+04 | 3.887E+03 | 6.574E+04 | 1.348E+05 |
| Ac20 | 79 | 1.2 | 2.281E+06 | 3.630E+04 | 1.033E+05 | 2.404E+05 | 2.504E+05 | 2.127E+05 | 2.448E+04 | 5.289E+05 | 8.848E+05 |
| Ac16 | 32 | 0.6 | 2.496E+06 | 2.846E+04 | 1.643E+05 | 2.433E+05 | 2.534E+05 | 2.514E+05 | 4.405E+04 | 4.724E+05 | 1.039E+06 |
| Ac16 | 40 | 0.6 | 3.319E+06 | 3.546E+04 | 1.629E+05 | 3.264E+05 | 3.678E+05 | 3.123E+05 | 3.913E+04 | 7.221E+05 | 1.353E+06 |
| Ac16 | 30 | 0.6 | 3.195E+06 | 4.541E+04 | 2.138E+05 | 2.885E+05 | 2.943E+05 | 3.340E+05 | 3.348E+04 | 7.012E+05 | 1.284E+06 |
| Ac16 | 34 | 0.6 | 3.567E+06 | 3.761E+04 | 2.909E+05 | 3.666E+05 | 4.026E+05 | 2.967E+05 | 6.494E+04 | 6.544E+05 | 1.453E+06 |
| Ac16 | 62 | 1.2 | 3.612E+06 | 7.502E+04 | 2.219E+05 | 2.877E+05 | 3.089E+05 | 3.611E+05 | 5.183E+04 | 7.228E+05 | 1.582E+06 |
| Ac21 | 56 | 0.6 | 3.894E+06 | 2.900E+04 | 2.496E+05 | 3.089E+05 | 3.409E+05 | 3.586E+05 | 4.143E+04 | 1.050E+06 | 1.515E+06 |
| Ac20 | 80 | 1.2 | 3.905E+06 | 7.521E+04 | 2.364E+05 | 3.560E+05 | 4.044E+05 | 4.400E+05 | 2.742E+04 | 7.471E+05 | 1.619E+06 |
| Ac16 | 42 | 0.6 | 3.832E+06 | 3.880E+04 | 2.902E+05 | 3.538E+05 | 4.397E+05 | 4.044E+05 | 5.770E+04 | 6.106E+05 | 1.637E+06 |
| Ac21 | 05 | 1.2 | 4.091E+06 | 6.181E+04 | 2.172E+05 | 3.556E+05 | 3.606E+05 | 3.792E+05 | 2.113E+04 | 9.713E+05 | 1.724E+06 |
| Ac21 | 54 | 0.6 | 4.211E+06 | 3.982E+04 | 2.440E+05 | 4.566E+05 | 4.620E+05 | 3.538E+05 | 4.110E+04 | 8.899E+05 | 1.724E+06 |
| Ac20 | 78 | 1.2 | 4.561E+06 | 9.814E+04 | 1.837E+05 | 4.196E+05 | 4.311E+05 | 6.263E+05 | 5.048E+04 | 1.218E+06 | 1.533E+06 |
| Ac21 | 010 | 1.2 | 4.735E+06 | 8.835E+04 | 3.978E+05 | 4.189E+05 | 4.540E+05 | 5.643E+05 | 3.624E+04 | 9.899E+05 | 1.785E+06 |
| Ac16 | 61 | 1.2 | 4.939E+06 | 1.090E+05 | 2.450E+05 | 5.484E+05 | 5.230E+05 | 4.321E+05 | 4.770E+04 | 8.238E+05 | 2.210E+06 |
| Ac16 | 46_19 | 1.2 | 5.728E+06 | 7.991E+04 | 3.777E+05 | 5.746E+05 | 5.316E+05 | 6.146E+05 | 6.868E+04 | 9.954E+05 | 2.485E+06 |
| Ac16 | 31_90 | 1.8 | 5.809E+06 | 1.383E+05 | 4.091E+05 | 5.220E+05 | 5.912E+05 | 7.216E+05 | 6.178E+04 | 1.251E+06 | 2.114E+06 |
| Ac16 | 34_90 | 1.8 | 5.890E+06 | 1.163E+05 | 2.623E+05 | 5.695E+05 | 5.223E+05 | 6.450E+05 | 8.889E+04 | 1.336E+06 | 2.350E+06 |
| Ac21 | 50 | 0.6 | 6.488E+06 | 6.230E+04 | 3.728E+05 | 5.445E+05 | 7.528E+05 | 5.869E+05 | 6.727E+04 | 1.274E+06 | 2.828E+06 |
| Ac22 | 19 | 2.4 | 6.366E+06 | 1.708E+05 | 4.770E+05 | 6.914E+05 | 6.302E+05 | 5.660E+05 | 5.224E+04 | 1.258E+06 | 2.520E+06 |
| Ac21 | 59 | 2.4 | 6.813E+06 | 1.921E+05 | 5.263E+05 | 6.560E+05 | 6.803E+05 | 9.307E+05 | 5.156E+04 | 1.031E+06 | 2.745E+06 |
| Ac16 | 24 | 2.4 | 7.328E+06 | 1.570E+05 | 5.994E+05 | 7.087E+05 | 7.443E+05 | 6.675E+05 | 6.154E+04 | 1.659E+06 | 2.730E+06 |
| Ac16 | 25 | 2.4 | 7.491E+06 | 1.718E+05 | 4.785E+05 | 6.926E+05 | 7.050E+05 | 9.608E+05 | 9.617E+04 | 1.777E+06 | 2.609E+06 |
| Ac16 | 42_90 | 1.8 | 7.646E+06 | 1.538E+05 | 5.703E+05 | 7.974E+05 | 8.343E+05 | 7.348E+05 | 8.166E+04 | 1.034E+06 | 3.440E+06 |
| Ac16 | 55 | 2.4 | 8.427E+06 | 2.614E+05 | 5.144E+05 | 8.561E+05 | 8.203E+05 | 9.793E+05 | 4.903E+04 | 1.447E+06 | 3.500E+06 |
| Ac21 | 52 | 3 | 9.011E+06 | 3.958E+05 | 6.311E+05 | 9.382E+05 | 9.846E+05 | 9.044E+05 | 1.425E+04 | 1.429E+06 | 3.713E+06 |
| Ac16 | 56 | 2.4 | 9.686E+06 | 2.009E+05 | 7.221E+05 | 8.496E+05 | 9.604E+05 | 9.527E+05 | 8.119E+04 | 2.473E+06 | 3.446E+06 |
| Ac09 | 97 | 3 | 1.113E+07 | 2.817E+05 | 8.021E+05 | 1.055E+06 | 1.089E+06 | 1.302E+06 | 1.525E+05 | 2.602E+06 | 3.844E+06 |
| Ac22 | 16 | 2.4 | 1.155E+07 | 2.803E+05 | 7.203E+05 | 1.016E+06 | 1.119E+06 | 1.173E+06 | 4.275E+04 | 2.309E+06 | 4.890E+06 |
| Ac21 | 15 | 3 | 1.260E+07 | 5.338E+05 | 7.714E+05 | 1.170E+06 | 1.290E+06 | 1.353E+06 | 9.966E+04 | 3.036E+06 | 4.343E+06 |
| Ac20 | 074 | 3 | 1.472E+07 | 4.931E+05 | 1.414E+06 | 1.316E+06 | 1.343E+06 | 1.276E+06 | 1.316E+05 | 3.092E+06 | 5.652E+06 |
| Ac21 | 38 | 3 | 1.508E+07 | 5.833E+05 | 9.186E+05 | 1.313E+06 | 1.376E+06 | 1.478E+06 | 9.015E+04 | 2.756E+06 | 6.563E+06 |
| Ac21 | 16 | 3 | 1.534E+07 | 6.886E+05 | 9.659E+05 | 1.312E+06 | 1.546E+06 | 1.266E+06 | 5.418E+04 | 4.395E+06 | 5.116E+06 |
| Ac21 | 19 | 3 | 1.641E+07 | 6.524E+05 | 8.672E+05 | 1.293E+06 | 1.424E+06 | 1.381E+06 | 8.315E+04 | 4.244E+06 | 6.467E+06 |
| Ac20 | 76 | 3 | 1.721E+07 | 8.682E+05 | 1.137E+06 | 1.711E+06 | 1.786E+06 | 1.840E+06 | 3.579E+04 | 3.502E+06 | 6.329E+06 |
| Ac16 | 44 | 0.6 | 1.875E+07 | 1.861E+05 | 9.291E+05 | 1.695E+06 | 1.927E+06 | 1.652E+06 | 2.816E+05 | 3.059E+06 | 9.036E+06 |
| Ac16 | 27 | 3 | 1.803E+07 | 6.541E+05 | 7.532E+05 | 1.497E+06 | 1.643E+06 | 1.831E+06 | 1.178E+05 | 4.612E+06 | 6.924E+06 |
| Ac22 | 15 | 2.4 | 3.062E+07 | 6.507E+05 | 2.447E+06 | 2.847E+06 | 3.030E+06 | 3.006E+06 | 1.503E+05 | 6.604E+06 | 1.188E+07 |
| Ac21 | 033 | 1.8 | 2.515E+07 | 6.934E+05 | 2.016E+06 | 2.439E+06 | 2.452E+06 | 3.298E+06 | 2.601E+05 | 5.167E+06 | 8.823E+06 |

**Supplementary Table 2.** Proportional volumes of brain compartments as percent of total brain tissue measured for each individual. Bin indicates HW (mm) category, divided into three groups (bins) for media workers. HW indicates actual size measurements of sampled individuals according to bin grouping.

| Colony | ID | Bin | OL | AL | MB-MC | MB-LC | MB-P | CB | SEG | ROCB | MBS |
| --- | --- | --- | --- | --- | --- | --- | --- | --- | --- | --- | --- |
| Ac16_30 | 30 | 0.6 | 0.014 | 0.067 | 0.090 | 0.092 | 0.105 | 0.010 | 0.219 | 0.402 | 0.287 |
| Ac16_32 | 32 | 0.6 | 0.011 | 0.066 | 0.097 | 0.102 | 0.101 | 0.018 | 0.189 | 0.416 | 0.300 |
| Ac16_34 | 34 | 0.6 | 0.011 | 0.082 | 0.103 | 0.113 | 0.083 | 0.018 | 0.183 | 0.407 | 0.299 |
| Ac16_40 | 40 | 0.6 | 0.011 | 0.049 | 0.098 | 0.111 | 0.094 | 0.012 | 0.218 | 0.408 | 0.303 |
| Ac16_42 | 42 | 0.6 | 0.010 | 0.076 | 0.092 | 0.115 | 0.106 | 0.015 | 0.159 | 0.427 | 0.313 |
| Ac16_44 | 44 | 0.6 | 0.010 | 0.050 | 0.090 | 0.103 | 0.088 | 0.015 | 0.163 | 0.482 | 0.281 |
| Ac20_006 | 006 | 0.6 | 0.013 | 0.056 | 0.078 | 0.089 | 0.111 | 0.017 | 0.151 | 0.484 | 0.279 |
| Ac21_50 | 50 | 0.6 | 0.010 | 0.057 | 0.084 | 0.116 | 0.090 | 0.010 | 0.196 | 0.436 | 0.290 |
| Ac21_54 | 54 | 0.6 | 0.009 | 0.058 | 0.108 | 0.110 | 0.084 | 0.010 | 0.211 | 0.409 | 0.302 |
| Ac21_56 | 56 | 0.6 | 0.007 | 0.064 | 0.079 | 0.088 | 0.092 | 0.011 | 0.270 | 0.389 | 0.259 |
| Ac16_46_19 | 46_19 | 1.2 | 0.014 | 0.066 | 0.100 | 0.093 | 0.107 | 0.012 | 0.174 | 0.434 | 0.300 |
| Ac16_61 | 61 | 1.2 | 0.022 | 0.050 | 0.111 | 0.106 | 0.087 | 0.010 | 0.167 | 0.447 | 0.304 |
| Ac16_62 | 62 | 1.2 | 0.021 | 0.061 | 0.080 | 0.086 | 0.100 | 0.014 | 0.200 | 0.438 | 0.265 |
| Ac20_055 | 055 | 1.2 | 0.014 | 0.067 | 0.098 | 0.105 | 0.127 | 0.014 | 0.203 | 0.371 | 0.330 |
| Ac20_056 | 056 | 1.2 | 0.020 | 0.069 | 0.102 | 0.101 | 0.132 | 0.013 | 0.194 | 0.368 | 0.335 |
| Ac20_78 | 78 | 1.2 | 0.022 | 0.040 | 0.092 | 0.095 | 0.137 | 0.011 | 0.267 | 0.336 | 0.324 |
| Ac20_79 | 79 | 1.2 | 0.016 | 0.045 | 0.105 | 0.110 | 0.093 | 0.011 | 0.232 | 0.388 | 0.308 |
| Ac20_80 | 80 | 1.2 | 0.019 | 0.061 | 0.091 | 0.104 | 0.113 | 0.007 | 0.191 | 0.415 | 0.307 |
| Ac21_010 | 010 | 1.2 | 0.019 | 0.084 | 0.088 | 0.096 | 0.119 | 0.008 | 0.209 | 0.377 | 0.304 |
| Ac21_05 | 05 | 1.2 | 0.015 | 0.053 | 0.087 | 0.088 | 0.093 | 0.005 | 0.237 | 0.421 | 0.268 |
| Ac16_31_90 | 31_90 | 1.8 | 0.024 | 0.070 | 0.090 | 0.102 | 0.124 | 0.011 | 0.215 | 0.364 | 0.316 |
| Ac16_34_90 | 34_90 | 1.8 | 0.020 | 0.045 | 0.097 | 0.089 | 0.110 | 0.015 | 0.227 | 0.399 | 0.295 |
| Ac16_42_90 | 42_90 | 1.8 | 0.020 | 0.075 | 0.104 | 0.109 | 0.096 | 0.011 | 0.135 | 0.450 | 0.310 |
| Ac20_033 | 033 | 1.8 | 0.020 | 0.088 | 0.105 | 0.105 | 0.104 | 0.008 | 0.162 | 0.410 | 0.314 |
| Ac20_036 | 036 | 1.8 | 0.025 | 0.077 | 0.103 | 0.099 | 0.142 | 0.009 | 0.148 | 0.398 | 0.344 |
| Ac20_29 | 29 | 1.8 | 0.022 | 0.061 | 0.104 | 0.100 | 0.116 | 0.007 | 0.154 | 0.435 | 0.320 |
| Ac20_34 | 34 | 1.8 | 0.023 | 0.078 | 0.097 | 0.097 | 0.093 | 0.008 | 0.211 | 0.393 | 0.287 |
| Ac21_033 | 033 | 1.8 | 0.028 | 0.080 | 0.097 | 0.098 | 0.131 | 0.010 | 0.205 | 0.351 | 0.326 |
| Ac21_30 | 30 | 1.8 | 0.021 | 0.064 | 0.100 | 0.098 | 0.132 | 0.015 | 0.177 | 0.393 | 0.330 |
| Ac21_36 | 36 | 1.8 | 0.029 | 0.068 | 0.097 | 0.101 | 0.115 | 0.014 | 0.173 | 0.403 | 0.314 |
| Ac16_24 | 24 | 2.4 | 0.021 | 0.082 | 0.097 | 0.102 | 0.091 | 0.008 | 0.226 | 0.373 | 0.289 |
| Ac16_25 | 25 | 2.4 | 0.023 | 0.064 | 0.092 | 0.094 | 0.128 | 0.013 | 0.237 | 0.348 | 0.315 |
| Ac16_55 | 55 | 2.4 | 0.031 | 0.061 | 0.102 | 0.097 | 0.116 | 0.006 | 0.172 | 0.415 | 0.315 |
| Ac16_56 | 56 | 2.4 | 0.021 | 0.075 | 0.088 | 0.099 | 0.098 | 0.008 | 0.255 | 0.356 | 0.285 |
| Ac20_21 | 21 | 2.4 | 0.031 | 0.097 | 0.093 | 0.102 | 0.109 | 0.005 | 0.176 | 0.387 | 0.304 |
| Ac21_28 | 28 | 2.4 | 0.027 | 0.091 | 0.093 | 0.097 | 0.118 | 0.011 | 0.185 | 0.378 | 0.308 |
| Ac21_59 | 59 | 2.4 | 0.028 | 0.077 | 0.096 | 0.100 | 0.137 | 0.008 | 0.151 | 0.403 | 0.333 |
| Ac22_15 | 15 | 2.4 | 0.021 | 0.080 | 0.093 | 0.099 | 0.098 | 0.005 | 0.216 | 0.388 | 0.290 |
| Ac22_16 | 16 | 2.4 | 0.024 | 0.062 | 0.088 | 0.097 | 0.102 | 0.004 | 0.200 | 0.423 | 0.286 |
| Ac22_19 | 19 | 2.4 | 0.027 | 0.075 | 0.109 | 0.099 | 0.089 | 0.008 | 0.198 | 0.396 | 0.297 |
| Ac09_97 | 97 | 3 | 0.025 | 0.072 | 0.095 | 0.098 | 0.117 | 0.014 | 0.234 | 0.345 | 0.310 |
| AC16_27 | 27 | 3 | 0.036 | 0.042 | 0.083 | 0.091 | 0.102 | 0.007 | 0.256 | 0.384 | 0.276 |
| Ac20_028 | 028 | 3 | 0.040 | 0.077 | 0.098 | 0.102 | 0.096 | 0.007 | 0.186 | 0.393 | 0.296 |
| Ac20_074 | 074 | 3 | 0.034 | 0.096 | 0.089 | 0.091 | 0.087 | 0.009 | 0.210 | 0.384 | 0.267 |
| Ac20_76 | 76 | 3 | 0.050 | 0.066 | 0.099 | 0.104 | 0.107 | 0.002 | 0.204 | 0.368 | 0.310 |
| Ac21_015 | 015 | 3 | 0.042 | 0.061 | 0.093 | 0.102 | 0.107 | 0.008 | 0.241 | 0.345 | 0.303 |
| Ac21_16 | 16 | 3 | 0.045 | 0.063 | 0.086 | 0.101 | 0.083 | 0.004 | 0.286 | 0.333 | 0.269 |
| Ac21_19 | 19 | 3 | 0.040 | 0.053 | 0.079 | 0.087 | 0.084 | 0.005 | 0.259 | 0.394 | 0.250 |
| Ac21_38 | 38 | 3 | 0.039 | 0.061 | 0.087 | 0.091 | 0.098 | 0.006 | 0.183 | 0.435 | 0.276 |
| Ac21_52 | 52 | 3 | 0.044 | 0.070 | 0.104 | 0.109 | 0.100 | 0.002 | 0.159 | 0.412 | 0.314 |

**Supplementary Table 3.**

Shapiro-Wilk test significance values used in determining whether data were normally distributed; significant p-value indicates non-normal distribution.

| Brain compartment | Absolute volume Shapiro-Wilk p value | Proportional volume Shapiro-Wilk p value |
| --- | --- | --- |
| Sum of neuropils | 2.201e-05 | N/A |
| Total neuropil volume/HW | 2.108e-09 | N/A |
| OL | 1.412e-07 | 0.017 |
| AL | 7.776e-07 | 0.822 |
| MB-S | 1.614e-05 | 0.810 |
| MB-MC | 2.446e-05 | 0.611 |
| MB-LC | 3.122e-05 | 0.518 |
| MB-P | 5.821e-06 | 0.032 |
| CX | 6.804e-07 | 0.822 |
| SEZ | 6.827e-06 | 0.593 |
| ROCB | 2.521e-05 | 0.597 |

**Supplementary Table 4.** Normalized *A. cephalotes* brain volume measurements (annotated by a different observer from that used to annotate brain images in main data set) used for linear discriminant analysis training. Bin indicates HW (mm) category, divided into three groups (bins) for media workers. HW indicates actual size measurements of sampled individuals according to bin grouping.

| Colony | ID | Worker | Bin | HW | OL | AL | MB-MC | MB-LC | MB-P | CB | SEG | ROCB |
| --- | --- | --- | --- | --- | --- | --- | --- | --- | --- | --- | --- | --- |
| Ac09 | 78 | Minim | 0.6 | 0.68 | 0.009 | 0.053 | 0.090 | 0.092 | 0.114 | 0.015 | 0.194 | 0.432 |
| Ac09 | 79 | Minim | 0.6 | 0.65 | 0.013 | 0.069 | 0.086 | 0.108 | 0.135 | 0.016 | 0.185 | 0.389 |
| Ac09 | 85 | Minim | 0.6 | 0.64 | 0.011 | 0.054 | 0.102 | 0.105 | 0.104 | 0.017 | 0.193 | 0.413 |
| Ac09 | 38 | Media | 1.2 | 1.25 | 0.019 | 0.082 | 0.092 | 0.085 | 0.090 | 0.011 | 0.205 | 0.416 |
| Ac09 | 44 | Media | 1.2 | 1.28 | 0.017 | 0.085 | 0.090 | 0.093 | 0.099 | 0.016 | 0.204 | 0.396 |
| Ac09 | 47 | Media | 1.2 | 1.13 | 0.013 | 0.081 | 0.091 | 0.090 | 0.119 | 0.014 | 0.190 | 0.402 |
| Ac09 | 50 | Media | 1.2 | 1.28 | 0.014 | 0.071 | 0.091 | 0.091 | 0.124 | 0.013 | 0.197 | 0.400 |
| Ac09 | 51 | Media | 1.2 | 1.13 | 0.016 | 0.074 | 0.100 | 0.094 | 0.113 | 0.013 | 0.166 | 0.425 |
| Ac09 | 53 | Media | 1.2 | 1.25 | 0.016 | 0.072 | 0.100 | 0.099 | 0.104 | 0.011 | 0.168 | 0.429 |
| Ac09 | 62 | Media | 1.2 | 1.2 | 0.016 | 0.067 | 0.095 | 0.094 | 0.131 | 0.016 | 0.184 | 0.397 |
| Ac09 | 1 | Media | 1.8 | 1.75 | 0.016 | 0.072 | 0.099 | 0.097 | 0.095 | 0.011 | 0.197 | 0.412 |
| Ac09 | 2 | Media | 1.8 | 1.78 | 0.022 | 0.090 | 0.087 | 0.082 | 0.094 | 0.006 | 0.231 | 0.388 |
| Ac09 | 5 | Media | 1.8 | 1.79 | 0.019 | 0.096 | 0.098 | 0.100 | 0.095 | 0.007 | 0.167 | 0.418 |
| Ac09 | 8 | Media | 1.8 | 1.78 | 0.021 | 0.072 | 0.106 | 0.104 | 0.104 | 0.007 | 0.210 | 0.376 |
| Ac09 | 12 | Media | 1.8 | 1.84 | 0.020 | 0.076 | 0.095 | 0.096 | 0.092 | 0.005 | 0.216 | 0.400 |
| Ac09 | 32 | Media | 1.8 | 1.73 | 0.017 | 0.066 | 0.090 | 0.092 | 0.117 | 0.011 | 0.215 | 0.392 |
| Ac09 | 33 | Media | 1.8 | 1.8 | 0.019 | 0.087 | 0.085 | 0.100 | 0.122 | 0.008 | 0.199 | 0.380 |
| Ac09 | 34 | Media | 1.8 | 1.75 | 0.023 | 0.068 | 0.102 | 0.102 | 0.110 | 0.015 | 0.196 | 0.383 |
| Ac09 | 43 | Media | 2.4 | 2.3 | 0.020 | 0.114 | 0.095 | 0.098 | 0.109 | 0.010 | 0.206 | 0.348 |
| Ac09 | 77 | Media | 2.4 | 2.35 | 0.023 | 0.074 | 0.106 | 0.092 | 0.095 | 0.011 | 0.158 | 0.442 |
| Ac09 | 81 | Media | 2.4 | 2.37 | 0.028 | 0.066 | 0.079 | 0.098 | 0.085 | 0.012 | 0.231 | 0.403 |
| Ac09 | 42 | Major | 3 | 3.9 | 0.033 | 0.079 | 0.103 | 0.093 | 0.091 | 0.008 | 0.191 | 0.401 |
| Ac09 | 68 | Major | 3 | 4.1 | 0.039 | 0.083 | 0.091 | 0.095 | 0.096 | 0.005 | 0.208 | 0.382 |
| Ac09 | 69 | Major | 3 | 3.9 | 0.042 | 0.100 | 0.106 | 0.108 | 0.097 | 0.005 | 0.147 | 0.395 |

**Supplementary Table 5.** Wilcoxon rank sum pairwise statistics for total brain volumes

**a.** Absolute total brain volume

| Worker size groups compared (HW, in mm) | Wilcoxon rank sum *post hoc* |
| --- | --- |
| 0.6-1.2 | 1.000 |
| 0.6-1.8 | 1.000 |
| 0.6-2.4 | 0.753 |
| 0.6-3 | 0.147 |
| 1.2-1.8 | 1.000 |
| 1.2-2.4 | 0.089 |
| 1.2-3 | 0.007 |
| 1.8-2.4 | 0.185 |
| 1.8-3 | 0.039 |
| 2.4-3 | 0.355 |

**b.** Total brain volume/individual HW

for each sample

| Worker size groups compared (HW, in mm) | Wilcoxon rank sum *post hoc* |
| --- | --- |
| 0.6-1.2 | 0.039 |
| 0.6-1.8 | 0.039 |
| 0.6-2.4 | 0.185 |
| 0.6-3 | 0.185 |
| 1.2-1.8 | 1.000 |
| 1.2-2.4 | 1.000 |
| 1.2-3 | 1.000 |
| 1.8-2.4 | 1.000 |
| 1.8-3 | 0.892 |
| 2.4-3 | 1.000 |

**Supplementary Table 6.** Wilcoxon rank sum pairwise statistics for absolute volumes

**a.** OL

| Worker size groups compared (HW, in mm) | Wilcoxon rank sum *post hoc* |
| --- | --- |
| 0.6-1.2 | 1.000 |
| 0.6-1.8 | 1.000 |
| 0.6-2.4 | 0.288 |
| 0.6-3 | 0.011 |
| 1.2-1.8 | 1.000 |
| 1.2-2.4 | 0.089 |
| 1.2-3 | 0.007 |
| 1.8-2.4 | 0.089 |
| 1.8-3 | 0.029 |
| 2.4-3 | 0.039 |

**b.** AL

| Worker size groups compared (HW, in mm) | Wilcoxon rank sum *post hoc* |
| --- | --- |
| 0.6-1.2 | 1.000 |
| 0.6-1.8 | 1.000 |
| 0.6-2.4 | 0.630 |
| 0.6-3 | 0.068 |
| 1.2-1.8 | 1.000 |
| 1.2-2.4 | 0.089 |
| 1.2-3 | 0.007 |
| 1.8-2.4 | 0.185 |
| 1.8-3 | 0.039 |
| 2.4-3 | 0.288 |

**c.** MB-S

| Worker size groups compared (HW, in mm) | Wilcoxon rank sum *post hoc* |
| --- | --- |
| 0.6-1.2 | 1.000 |
| 0.6-1.8 | 1.000 |
| 0.6-2.4 | 0.630 |
| 0.6-3 | 0.115 |
| 1.2-1.8 | 1.000 |
| 1.2-2.4 | 0.089 |
| 1.2-3 | 0.007 |
| 1.8-2.4 | 0.185 |
| 1.8-3 | 0.052 |
| 2.4-3 | 0.232 |

**d.** MB-MC

| Worker size groups compared (HW, in mm) | Wilcoxon rank sum *post hoc* |
| --- | --- |
| 0.6-1.2 | 1.000 |
| 0.6-1.8 | 1.000 |
| 0.6-2.4 | 0.630 |
| 0.6-3 | 0.115 |
| 1.2-1.8 | 1.000 |
| 1.2-2.4 | 0.089 |
| 1.2-3 | 0.007 |
| 1.8-2.4 | 0.185 |
| 1.8-3 | 0.052 |
| 2.4-3 | 0.232 |

**e.** MB-LC

| Worker size groups compared (HW, in mm) | Wilcoxon rank sum *post hoc* |
| --- | --- |
| 0.6-1.2 | 1.000 |
| 0.6-1.8 | 1.000 |
| 0.6-2.4 | 1.000 |
| 0.6-3 | 0.147 |
| 1.2-1.8 | 1.000 |
| 1.2-2.4 | 0.089 |
| 1.2-3 | 0.007 |
| 1.8-2.4 | 0.232 |
| 1.8-3 | 0.039 |
| 2.4-3 | 0.288 |

**f.** MB-P

| Worker size groups compared (HW, in mm) | Wilcoxon rank sum *post hoc* |
| --- | --- |
| 0.6-1.2 | 1.000 |
| 0.6-1.8 | 1.000 |
| 0.6-2.4 | 0.753 |
| 0.6-3 | 0.089 |
| 1.2-1.8 | 1.000 |
| 1.2-2.4 | 0.147 |
| 1.2-3 | 0.007 |
| 1.8-2.4 | 0.355 |
| 1.8-3 | 0.115 |
| 2.4-3 | 0.524 |

**g.** CX

| Worker size groups compared (HW, in mm) | Wilcoxon rank sum *post hoc* |
| --- | --- |
| 0.6-1.2 | 1.000 |
| 0.6-1.8 | 1.000 |
| 0.6-2.4 | 1.000 |
| 0.6-3 | 1.000 |
| 1.2-1.8 | 1.000 |
| 1.2-2.4 | 1.000 |
| 1.2-3 | 0.520 |
| 1.8-2.4 | 1.000 |
| 1.8-3 | 0.630 |
| 2.4-3 | 1.000 |

**h.** SEZ

| Worker size groups compared (HW, in mm) | Wilcoxon rank sum *post hoc* |
| --- | --- |
| 0.6-1.2 | 1.000 |
| 0.6-1.8 | 1.000 |
| 0.6-2.4 | 1.000 |
| 0.6-3 | 0.039 |
| 1.2-1.8 | 1.000 |
| 1.2-2.4 | 0.115 |
| 1.2-3 | 0.007 |
| 1.8-2.4 | 0.019 |
| 1.8-3 | 0.039 |
| 2.4-3 | 0.524 |

**i.** ROCB

| Worker size groups compared (HW, in mm) | Wilcoxon rank sum *post hoc* |
| --- | --- |
| 0.6-1.2 | 1.000 |
| 0.6-1.8 | 1.000 |
| 0.6-2.4 | 1.000 |
| 0.6-3 | 0.147 |
| 1.2-1.8 | 1.000 |
| 1.2-2.4 | 0.089 |
| 1.2-3 | 0.007 |
| 1.8-2.4 | 0.185 |
| 1.8-3 | 0.039 |
| 2.4-3 | 0.355 |

**Supplementary Table 7.** Tukey and Wilcoxon rank sum pairwise statistics for relative volumes

**a.** OL

| Worker size groups compared (HW, in mm) | Wilcoxon rank sum *post hoc* |
| --- | --- |
| 0.6-1.2 | 2.000e-4 |
| 0.6-1.8 | 1.000e-4 |
| 0.6-2.4 | 1.000e-4 |
| 0.6-3 | 1.000e-4 |
| 1.2-1.8 | 0.068 |
| 1.2-2.4 | 0.005 |
| 1.2-3 | 1.000e-4 |
| 1.8-2.4 | 1.000 |
| 1.8-3 | 4.000e-4 |
| 2.4-3 | 0.002 |

**b.** AL

| Worker size groups compared (HW, in mm) | Tukey's *post hoc* |
| --- | --- |
| 0.6-1.2 | 0.988 |
| 0.6-1.8 | 0.585 |
| 0.6-2.4 | 0.108 |
| 0.6-3 | 0.962 |
| 1.2-1.8 | 0.300 |
| 1.2-2.4 | 0.034 |
| 1.2-3 | 0.772 |
| 1.8-2.4 | 0.841 |
| 1.8-3 | 0.929 |
| 2.4-3 | 0.370 |

**c.** MB-S

| Worker size groups compared (HW, in mm) | Tukey’s *post hoc* |
| --- | --- |
| 0.6-1.2 | 0.517 |
| 0.6-1.8 | 0.048 |
| 0.6-2.4 | 0.696 |
| 0.6-3 | 0.987 |
| 1.2-1.8 | 0.707 |
| 1.2-2.4 | 0.998 |
| 1.2-3 | 0.245 |
| 1.8-2.4 | 0.529 |
| 1.8-3 | 0.013 |
| 2.4-3 | 0.387 |

**d.** MB-MC

| Worker size groups compared (HW, in mm) | Tukey's *post hoc* |
| --- | --- |
| 0.6-1.2 | 0.876 |
| 0.6-1.8 | 0.270 |
| 0.6-2.4 | 0.928 |
| 0.6-3 | 0.999 |
| 1.2-1.8 | 0.817 |
| 1.2-2.4 | 1.000 |
| 1.2-3 | 0.752 |
| 1.8-2.4 | 0.738 |
| 1.8-3 | 0.171 |
| 2.4-3 | 0.829 |

**e.** MB-LC

| Worker size groups compared (HW, in mm) | Tukey's *post hoc* |
| --- | --- |
| 0.6-1.2 | 0.452 |
| 0.6-1.8 | 0.748 |
| 0.6-2.4 | 0.529 |
| 0.6-3 | 0.346 |
| 1.2-1.8 | 0.989 |
| 1.2-2.4 | 1.000 |
| 1.2-3 | 1.000 |
| 1.8-2.4 | 0.996 |
| 1.8-3 | 0.962 |
| 2.4-3 | 0.998 |

**f.** MB-P

| Worker size groups compared (HW, in mm) | Wilcoxon rank sum *post hoc* |
| --- | --- |
| 0.6-1.2 | 0.433 |
| 0.6-1.8 | 0.039 |
| 0.6-2.4 | 0.630 |
| 0.6-3 | 1.000 |
| 1.2-1.8 | 1.000 |
| 1.2-2.4 | 1.000 |
| 1.2-3 | 1.000 |
| 1.8-2.4 | 1.000 |
| 1.8-3 | 0.232 |
| 2.4-3 | 1.000 |

**g.** CX

| Worker size groups compared (HW, in mm) | Tukey's *post hoc* |
| --- | --- |
| 0.6-1.2 | 0.212 |
| 0.6-1.8 | 0.265 |
| 0.6-2.4 | 0.001 |
| 0.6-3 | 4.830e-05 |
| 1.2-1.8 | 1.000 |
| 1.2-2.4 | 0.238 |
| 1.2-3 | 0.033 |
| 1.8-2.4 | 0.189 |
| 1.8-3 | 0.024 |
| 2.4-3 | 0.890 |

**h.** SEZ

| Worker size groups compared (HW, in mm) | Tukey's *post hoc* |
| --- | --- |
| 0.6-1.2 | 0.945 |
| 0.6-1.8 | 0.854 |
| 0.6-2.4 | 0.996 |
| 0.6-3 | 0.462 |
| 1.2-1.8 | 0.419 |
| 1.2-2.4 | 0.995 |
| 1.2-3 | 0.885 |
| 1.8-2.4 | 0.658 |
| 1.8-3 | 0.075 |
| 2.4-3 | 0.684 |

**i.** ROCB

| Worker size groups compared (HW, in mm) | Tukey's *post hoc* |
| --- | --- |
| 0.6-1.2 | 0.332 |
| 0.6-1.8 | 0.330 |
| 0.6-2.4 | 0.054 |
| 0.6-3 | 0.014 |
| 1.2-1.8 | 1.000 |
| 1.2-2.4 | 0.892 |
| 1.2-3 | 0.607 |
| 1.8-2.4 | 0.893 |
| 1.8-3 | 0.609 |
| 2.4-3 | 0.984 |
